# Modular Architecture of the SAGA Complex Governs Stress Adaptation, Morphogenesis, and Histone Acetylation in *Candida albicans*

**DOI:** 10.64898/2026.02.06.704420

**Authors:** Poonam Poonia, Manjit Kumar Srivastav, Priyanka Nagar, Krishnamurthy Natarajan

## Abstract

The SAGA complex is a conserved, multifunctional transcriptional co-activator known for its roles in chromatin modification and transcriptional regulation. While SAGA has been extensively characterized in *Saccharomyces cerevisiae* and metazoans, its modular organization and functional significance in the major human fungal pathogen *Candida albicans* remain poorly understood. Through bioinformatic analyses, we found that SAGA subunits are conserved in *C. albicans*. Genetic disruption of the histone acetyltransferase (HAT; *GCN5, ADA2*), structural (SPT; *SPT7, SPT20, TAF12L*), and TATA-binding protein interaction (TBP-interaction; *SPT3, SPT8*) modules leads to impaired growth under oxidative, metal, and antifungal stress conditions and causes severe defects in filamentation. In contrast, deletion of the deubiquitination (DUB) module components *UBP8* and *SUS1* results in minimal phenotypic consequences. Strikingly, loss of *SGF73*, a structural component linking the DUB module to the SAGA core, produces pronounced defects in stress conditions and filamentation, phenocopying *SPT3* and *SPT8* mutants. Consistent with these observations, filamentation-associated genes are significantly upregulated in *SGF73*, *SPT3* and *SPT8* mutants. Notably, these mutants also exhibit elevated global levels of histone H3 lysine-9 acetylation (H3K9ac), suggesting a critical role for *SGF73*-mediated SAGA integrity in coordinating chromatin acetylation with transcriptional programs governing stress responses and filamentation in *C. albicans*.

## Introduction

The SAGA (Spt–Ada–Gcn5 Acetyltransferase) complex is a conserved, multi-subunit transcriptional co-activator present from yeast to humans (1,2). Initially identified through genetic screens for suppressors of transcriptional defects in *Saccharomyces cerevisiae*, SAGA has since emerged as a central integrator of chromatin modification, transcription factor signaling, and RNA polymerase II (RNAPII) recruitment {Grant, 1997 #20}. The complex comprises 19 core subunits, along with additional species-specific components, and is organized into distinct yet functionally interdependent modules: the HAT (histone acetyltransferase) module (*GCN5, ADA2, ADA3, SGF29*), the core scaffold module (*SPT7, SPT20, ADA1*, and five TAFs), the TBP-interaction module (*SPT3, SPT8*), and the activator-interaction module (*TRA1)* (3–7). High-resolution cryo-EM studies have revealed that these modules form a dynamic architecture that undergoes conformational rearrangements upon activator binding, thereby enabling promoter-specific regulatory outcomes (3,5,8). Functionally, SAGA regulates RNAPII-dependent transcription with a prominent role at inducible, stress-responsive, and developmentally regulated genes (9,10). Genome-wide studies in yeast and humans have shown that SAGA recruits to most RNAPII promoters, with coordinated action among subunits essential for full transcriptional activity and enzymatic function (11,12). While individual modules contribute to distinct regulatory outcomes, their interdependence ensures complex integrity and gene-specific responsiveness (13,14).

*Candida albicans*, a major fungal pathogen, causes both superficial and life-threatening systemic infections, particularly in immunocompromised individuals (15–17). A defining feature of *C. albicans* pathogenesis is its remarkable transcriptional plasticity, which enables rapid adaptation to host-imposed stresses, including oxidative burst, nutrient limitation, metal sequestration, antifungal exposure, and immune surveillance (18,19). Central virulence traits, including reversible yeast-to-hypha transitions, biofilm formation, metabolic flexibility, and antifungal drug resistance, are tightly controlled by large transcriptional networks involving sequence-specific transcription factors, chromatin modifiers, and co-activator complexes (20). Although chromatin regulation has emerged as a key determinant of fungal virulence, our understanding of how conserved transcriptional co-activators operate in pathogenic fungi remains incomplete. Prior studies in *C. albicans* have implicated individual SAGA subunits, including Gcn5, Ada2, Tra1, and Spt3, in regulating filamentation, stress responses, and antifungal tolerance (21–28). However, these studies have largely focused on single subunits or limited subsets within specific modules. Consequently, how the modular organization of SAGA and the functional interplay between its modules collectively shape transcriptional regulation in *C. albicans* remains poorly defined.

Here, we present a systematic genetic dissection of the SAGA complex in *C. albicans*. Our findings demonstrate that SAGA is structurally conserved yet functionally diversified, operating in a module-specific manner to regulate transcriptional responses to environmental stress. Notably, we uncover antagonistic interactions between modules, with the HAT and TBP-interaction modules exerting opposing effects on filamentation and global histone acetylation. In contrast, the HAT and core scaffold modules differentially modulate sensitivity to the antifungal drug fluconazole. Moreover, we identify a previously unrecognized, non-canonical role for the DUB-module subunit *SGF73* in regulating filamentation and histone acetylation. Together, our findings reveal a modular and context-dependent regulatory framework through which the SAGA complex shapes transcriptional plasticity and pathogenic traits in *C. albicans*.

## Results

### SAGA subunits in *Candida albicans* retain conserved, functionally distinct domains

The diverse functions of the SAGA complex are carried out by dedicated subunits, each containing one or more evolutionarily conserved protein domains that enable their specialized roles. To investigate the evolutionary and functional conservation of the SAGA complex in *Candida albicans*, we first identified the putative orthologs of *S. cerevisiae* SAGA subunits in *C. albicans* using BLASTP analysis (genome assembly 22). Each of the 19 bona fide *S. cerevisiae* SAGA subunit protein sequences was used as a query against the *C. albicans* proteome. For all 19 subunits, we identified a single best-hit homolog with high sequence similarity and strong E-values, indicating a high degree of conservation (**Figure 1, Table 1**). We also recently reported the association of all these subunits by biochemical purification of the *Ca*SAGA complex (29). To further assess functional domain conservation between SAGA complex components, the amino acid sequences of the identified *C. albicans* SAGA subunits were analyzed using the SMART protein domain prediction tool (30). This analysis revealed that all previously characterized functional domains from *S. cerevisiae* SAGA subunits are also present in their *C. albicans* counterparts (**Figure 1**), further supporting evolutionary conservation of the *C. albicans* SAGA complex.

**Fig. 1.**
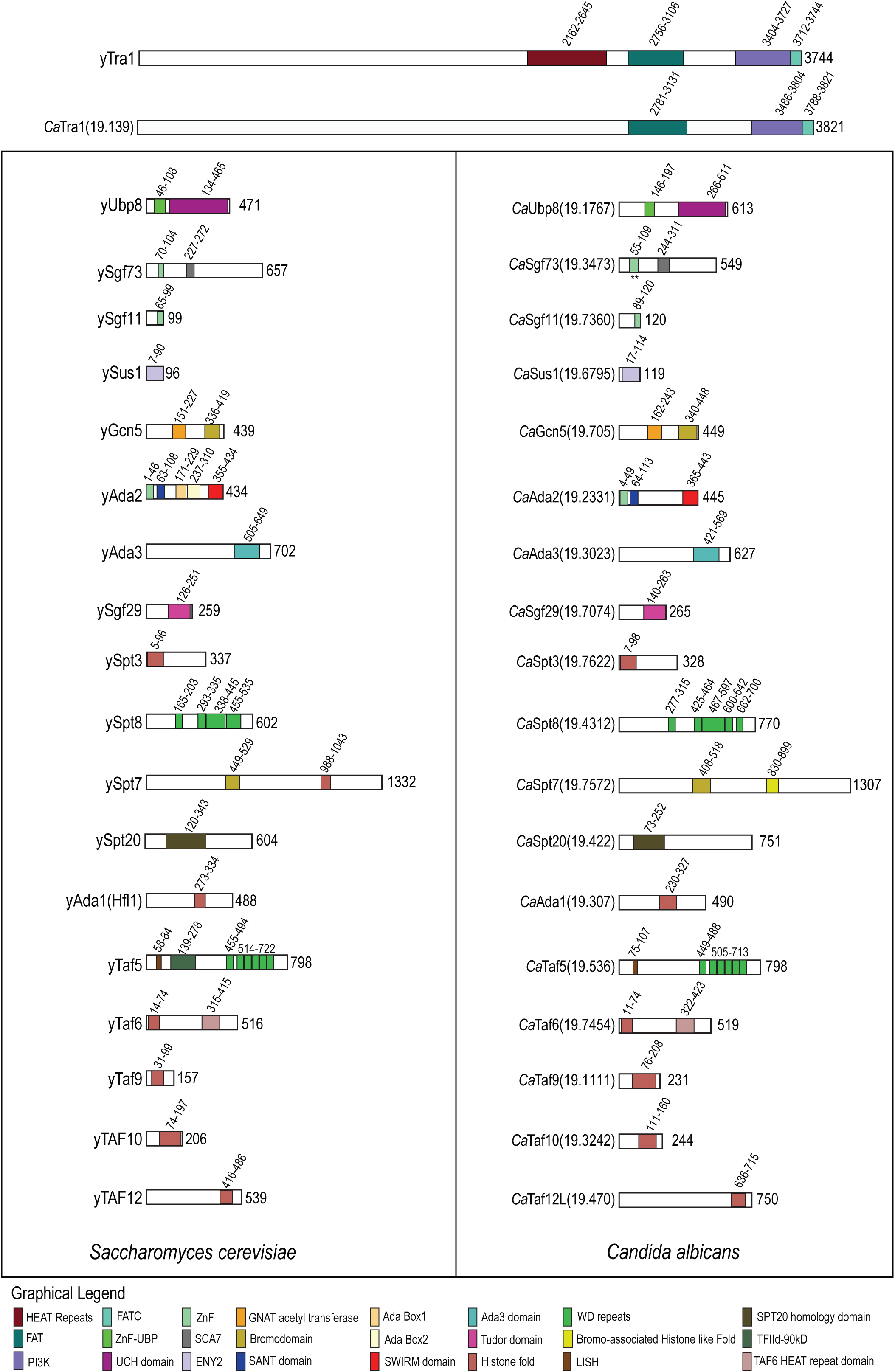
Cross-species comparison of the composition and domain architecture of SAGA subunits. Schematic diagrams of SAGA complex subunits from *Saccharomyces cerevisiae* and *Candida albicans* highlight evolutionary conservation and divergence in domain architecture. Known protein domains in *S. cerevisiae* were assigned according to (69). Putative domains in *C. albicans* subunits were predicted using SMART domain analysis, validated by sequence alignment with orthologous *S. cerevisiae* proteins, and supported by structural modeling using Phyre2. Amino acid coordinates indicate domain boundaries. Prefixes “y” and “Ca” denote yeast and *Candida* proteins, respectively. A graphical key defines the domain types represented. This analysis provides a foundation for understanding functional conservation and species-specific adaptations within the fungal SAGA complex.

To gain a broader perspective on the conservation of the SAGA complex within the fungal kingdom, we compared SAGA composition across several fungal species using published both *in silico* analyses and available biochemical purification data (**Table S1**). This comparative analysis showed that, although most SAGA subunits are conserved across diverse fungi, the DUB module exhibited the most variation. Notably, *C. albicans* (this study and (29)) and *Schizosaccharomyces pombe* (31) possess all the DUB module subunits *UBP8, SGF11, SGF73,* and *SUS1,* a composition similar to *S. cerevisiae*. In contrast, fungi such as *Cryptococcus neoformans* (32) and *Fusarium fujikuroi* (33) appear to lack one or more subunits of a functional DUB module. Notably, although *the Aspergillus nidulans* genome encodes all DUB module subunits, these do not co-purify with the remainder of the SAGA complex (34). These studies thus highlight a potential evolutionary divergence in the regulatory function of the DUB module among pathogenic and filamentous fungi.

### Distinct SAGA modules variably impact *Candida albicans* viability and fitness

To investigate the functional conservation of the modular architecture of SAGA complex in *C. albicans,* we constructed null mutant strains for nine representative subunits; *GCN5, ADA2* (HAT module); *SPT7, SPT20* (Core-module), *SPT3, SPT8* (TBP-interaction module) and *UBP8, SGF73* and *SUS1* (Deubiquitination module) spanning four functional modules of the *Ca*SAGA (Figure 2A). Viable null mutants were obtained for all these subunits, while in contrast, despite multiple deletion strategies and repeated attempts, we could not obtain null mutants for *ADA1* (core-module) and *TRA1* (TF-binding module), suggesting that these subunits may be essential for *C. albicans* viability. *TRA1* essentiality for *C. albicans* viability is highly probable, as it is essential in most organisms lacking a *TRA1* paralog, unlike *S. pombe*, which encodes a functionally specialized paralog, *TRA2* (31) (35). Similarly, we earlier showed that the viability of the *taf12lΔ/Δ* mutant reflects functional divergence between SAGA- and TFIID-associated TAF12 paralogs (26). While *ADA1* is nonessential in *S. cerevisiae* and *C. neoformans* (36), a study reported it essential in *C. glabrata* (37), suggesting possible *Candida*-specific roles.

**Fig. 2.**
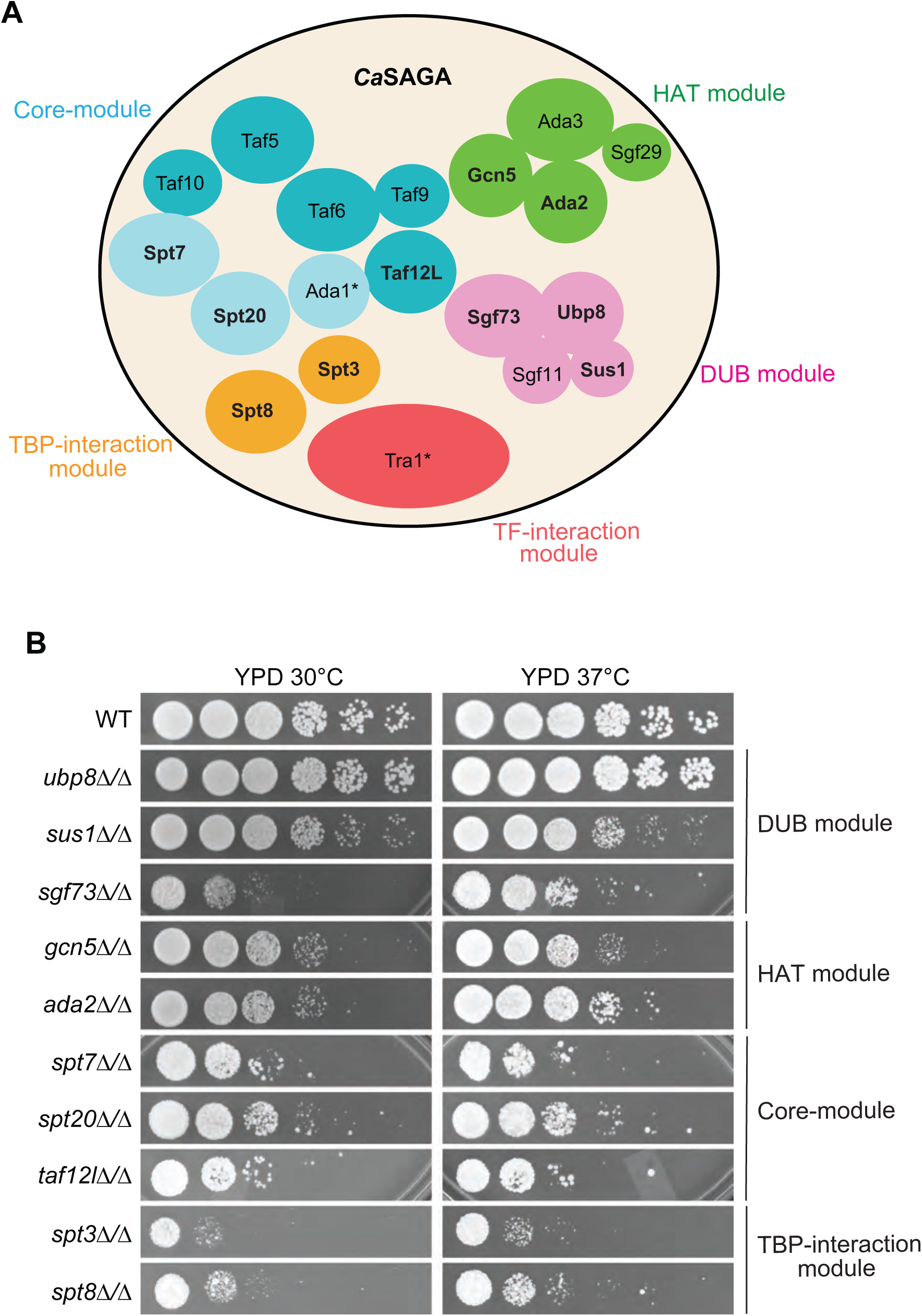
Deletion of SAGA subunits impairs the growth and fitness of *Candida albicans*. (A) Modular map of the *C. albicans* SAGA complex showing the positions of deleted subunits across the functional modules. (B) Growth phenotypes of SAGA deletion mutants were assessed by spot assays. Strains were cultured in YPD medium to saturation, serially diluted, and spotted onto YPD plates. Plates were incubated at 30°C and 37°C for up to 48 hours and imaged every 12 hours.

Next, we assessed the growth phenotypes of these null mutants at the optimal temperature (30°C) and under host environmental conditions (37°C). We also included only viable TAF mutant *(taf12lΔ/Δ)* in our analysis, which is required for virulence (26). With the exception of the *ubp8Δ/Δ* mutant (encoding the enzymatic subunit of the DUB module), all other deletion mutants displayed growth defects compared to the wild-type strain at both temperatures (**Figure 2B**). The severity of these growth defects varied considerably among different mutants and modules. Interestingly, module-specific growth patterns were observed. The HAT module mutants (*gcn5Δ/Δ* and *ada2Δ/*Δ) exhibited mild growth reduction (Figure 2B). In contrast, the core-module and TBP-interaction module mutants (*spt7Δ/Δ, spt20Δ/Δ,* and *spt8Δ/Δ spt3Δ/Δ,* respectively) showed severe growth impairment, highlighting the critical role of this module in cellular fitness. The TAF mutant part of the core module (*taf12lΔ/Δ*) also showed a severe growth phenotype similar to that of the SPT subunits of the core module (**Figure 2B**). Among the DUB module mutants, *sus1Δ/Δ* showed a moderate growth defect, whereas *sgf73Δ/Δ* exhibited a strong defect, comparable to the core-module mutants (**Figure 2B**).

Together, these findings demonstrate that while most individual *Ca*SAGA subunits (excluding *ADA1* and *TRA1*) are not essential for viability, their absence significantly impacts fungal growth. Moreover, the module-specific phenotypes suggest distinct contributions of each functional unit to the overall activity and structural integrity of the SAGA complex in *C. albicans*.

### Stress phenotypes of SAGA mutants are module-specific and evolutionarily conserved

During host colonization and infection, *Candida albicans* encounters a broad range of environmental stresses. The SAGA complex is a well-characterized regulator of stress responses in eukaryotes. To investigate the role of the SAGA complex in environmental stress adaptation in *C. albicans*, we analyzed the growth of strains lacking individual SAGA subunits under various stress conditions, including oxidative stress (menadione and hydrogen peroxide), heavy metal stress (cadmium chloride), amino acid starvation (sulfometuron methyl, SM), iron deprivation (4,7-diphenyl-1,10-phenanthrolinedisulfonic acid, BPS), and inhibition of transcription elongation (6-azauracil) (**Figure 3A-B**). Additionally, we assessed the sensitivity of these mutants to the antifungal drug fluconazole. Growth was scored semi-quantitatively on a 0–5 scale relative to the wild-type strain (**Figure 3C**). Mutants lacking subunits of the HAT module (*gcn5Δ/Δ*, *ada2Δ/Δ*), core and TBP-interaction modules (*spt7Δ/Δ*, *spt20Δ/Δ*, *taf12lΔ/Δ*, *spt3Δ/Δ*, *spt8Δ/Δ*), and *sgf73Δ/Δ* exhibited pronounced growth defects under multiple stress conditions (**Figure 3A–C**). Notably, except for the DUB module, mutants from the same functional module tended to exhibit similar phenotypes, suggesting modular coherence in stress response functions.

**Fig. 3.**
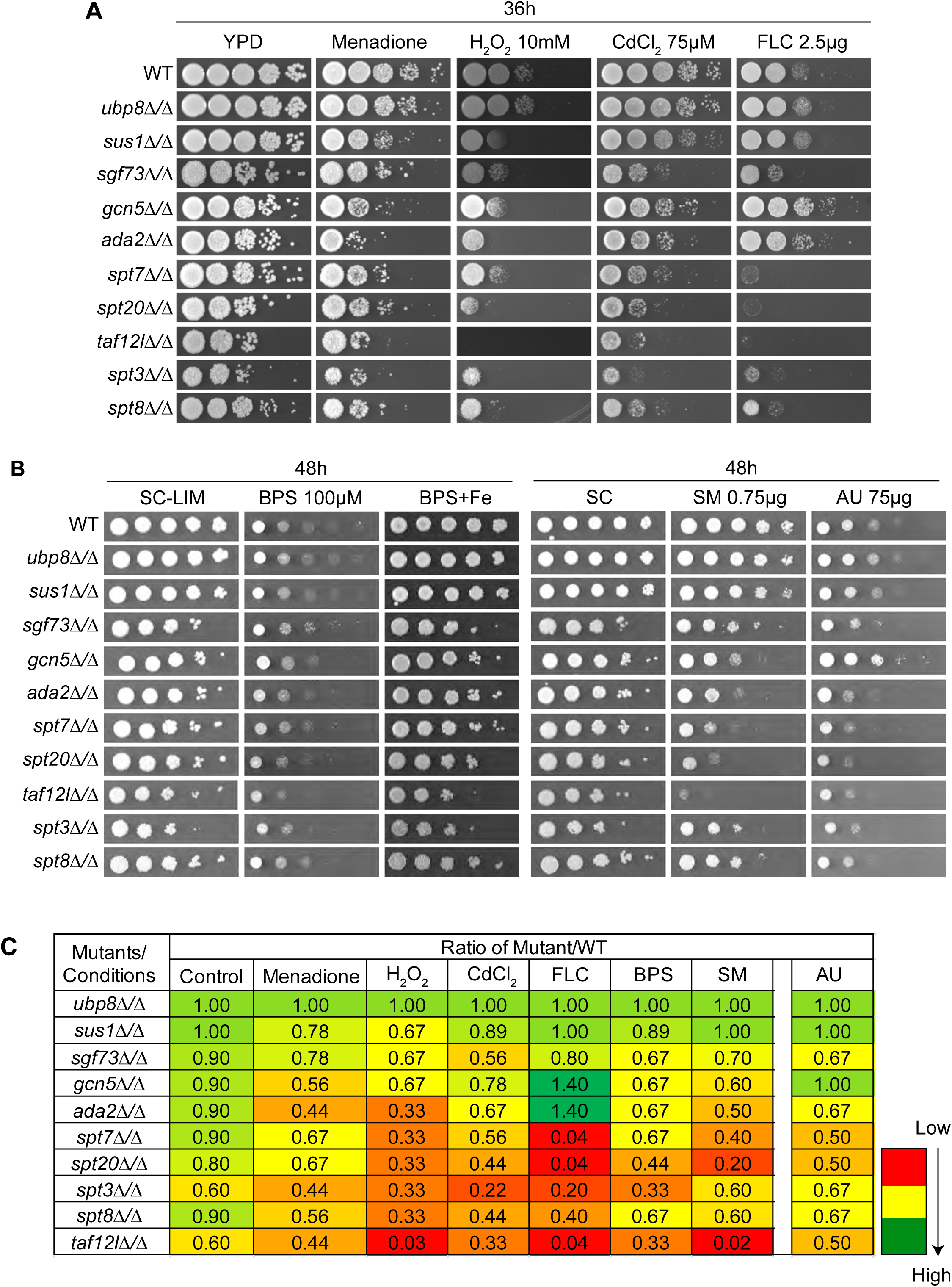
SAGA complex mediates stress adaptation in *C. albicans.* Strains were cultured in YPD medium to saturation, serially diluted, and spotted onto specified plates. Plates were incubated at 30°C for up to 48 hours and imaged every 12 hours. (A) Growth phenotype of different *Ca*SAGA deletion mutant strains under oxidative stress (90µM Menadione and 10 mM Hydrogen peroxide), metal stress (75µM Cadmium chloride), and fluconazole (2.5µg/ml) conditions. (B) Under iron limitation conditions, SC-LIM, SC-LIM+BPS (100µM), and SC-LIM+BPS+FAS (100µM). (C) Under amino acid starvation (SM 0.75µg/ml) and under 6-Azauracil (75µg/ml) conditions. (D) A colour-coded heatmap summarizes quantitative growth scores of SAGA deletion mutants. The strains were scored on a 0-5 scale based on growth across dilutions. The scores are represented as a ratio of mutant to WT for a particular condition and colour coded in Microsoft Excel. Control represents the scores for control media for different conditions: YPD for panel A, SC-LIM for panel B, and SC for panel C.

Of the ten mutants tested, all except *ubp8Δ/Δ* were sensitive to oxidative stress induced by both menadione and hydrogen peroxide (**Figure 3A**). While mutants within a module generally showed consistent stress sensitivities, we observed stressor-specific differences. For example, HAT module mutants (*gcn5Δ/Δ*, *ada2Δ/Δ*) showed heightened sensitivity to menadione compared to the core-module mutants (*spt7Δ/Δ*, *spt20Δ/Δ*, *spt3Δ/Δ*, *spt8Δ/Δ*), whereas all mutants displayed similar sensitivities to hydrogen peroxide. These findings underscore the importance of multiple SAGA activities, acetylation, deubiquitination, and TBP recruitment in orchestrating oxidative stress resistance. Under cadmium-induced heavy metal stress, deletion of core module subunits and *sgf73* resulted in severe growth defects, while the HAT module mutants (*gcn5Δ/Δ*, *ada2Δ/Δ*) and DUB subunit mutant *sus1Δ/Δ* exhibited moderate sensitivity (**Figure 3A**). In contrast, under iron deprivation and amino acid starvation, only *taf12lΔ/Δ* showed severe sensitivity, while other mutants displayed mild to no phenotypes. Similar trends were observed in response to the transcription elongation inhibitor 6-azauracil, with most mutants exhibiting moderate growth impairment (**Figure 3B**).

Exposure to fluconazole revealed divergent phenotypes across the SAGA mutants. Deletion of core (*spt7Δ/Δ*, *spt20Δ/Δ*, *taf12lΔ/Δ*) and TBP-interaction module (*spt3Δ/Δ*, *spt8Δ/Δ*) subunits, as well as *sgf73Δ/Δ*, resulted in increased drug sensitivity. Conversely, deletion of the HAT module components (*gcn5Δ/Δ*, *ada2Δ/Δ*) led to increased resistance to fluconazole. The DUB subunit mutants *ubp8Δ/Δ* and *sus1Δ/Δ* showed either no change or a slight improvement in growth in the presence of fluconazole (**Figure 3A**). Given the inherent genome instability of *C. albicans*, which can result in chromosomal aneuploidy or secondary genomic alterations, we performed complementation assays to confirm that the observed stress phenotypes were directly attributable to gene deletion. Reintroduction of the respective wild-type gene into *gcn5Δ/Δ*, *ada2Δ/Δ*, *spt7Δ/Δ*, *spt20Δ/Δ*, *taf12lΔ/Δ*, *spt3Δ/Δ*, *spt8Δ/Δ*, *sus1Δ/Δ*, and *sgf73Δ/Δ* restored growth under oxidative stress (menadione), fluconazole, and iron-deprivation conditions, supporting the conclusion that these phenotypes represent primary consequences of the gene deletions (**Figure S1**, (26)). To determine whether these stress-associated phenotypes are conserved across species, we performed parallel analyses in *Saccharomyces cerevisiae* strains carrying deletions of orthologous SAGA subunits (**Figure S2A–E**). The HAT and SPT module mutants in *S. cerevisiae* exhibited similar sensitivity profiles, including susceptibility to iron deprivation, a phenotype not previously linked to SAGA in this organism. As observed in *C. albicans*, deletion of the DUB subunit *UBP8* in *S. cerevisiae* did not result in detectable growth defects under any of the tested conditions.

Together, these findings highlight the modular and partially conserved role of the SAGA complex in mediating diverse stress responses in fungi. While the core functions of the HAT and SPT modules appear broadly conserved, species-specific adaptations, such as the opposing fluconazole phenotypes in *C. albicans,* suggest evolutionary rewiring of SAGA functionality to support the unique pathogenic strategies of this opportunistic fungus.

### The SAGA complex regulates morphogenesis and the transition to the hyphal form

The morphological transition from yeast to hyphal form is a key feature of *Candida albicans*, and this dynamic process has been closely linked to fungal pathogenesis (38) (39) (40). Previous studies have demonstrated that deletion of the individual SAGA subunit results in varied morphological phenotypes (21,23,26). However, it is unclear whether the SAGA complex functions as a whole or whether a specific SAGA module or activity is associated with particular morphological phenotypes. To investigate the complex, specific, and module-specific contributions of the SAGA complex to morphogenetic regulation, we analyzed cellular and colony morphology across our panel of SAGA subunit deletion mutants (**Figures 4** and **S3**).

**Fig. 4.**
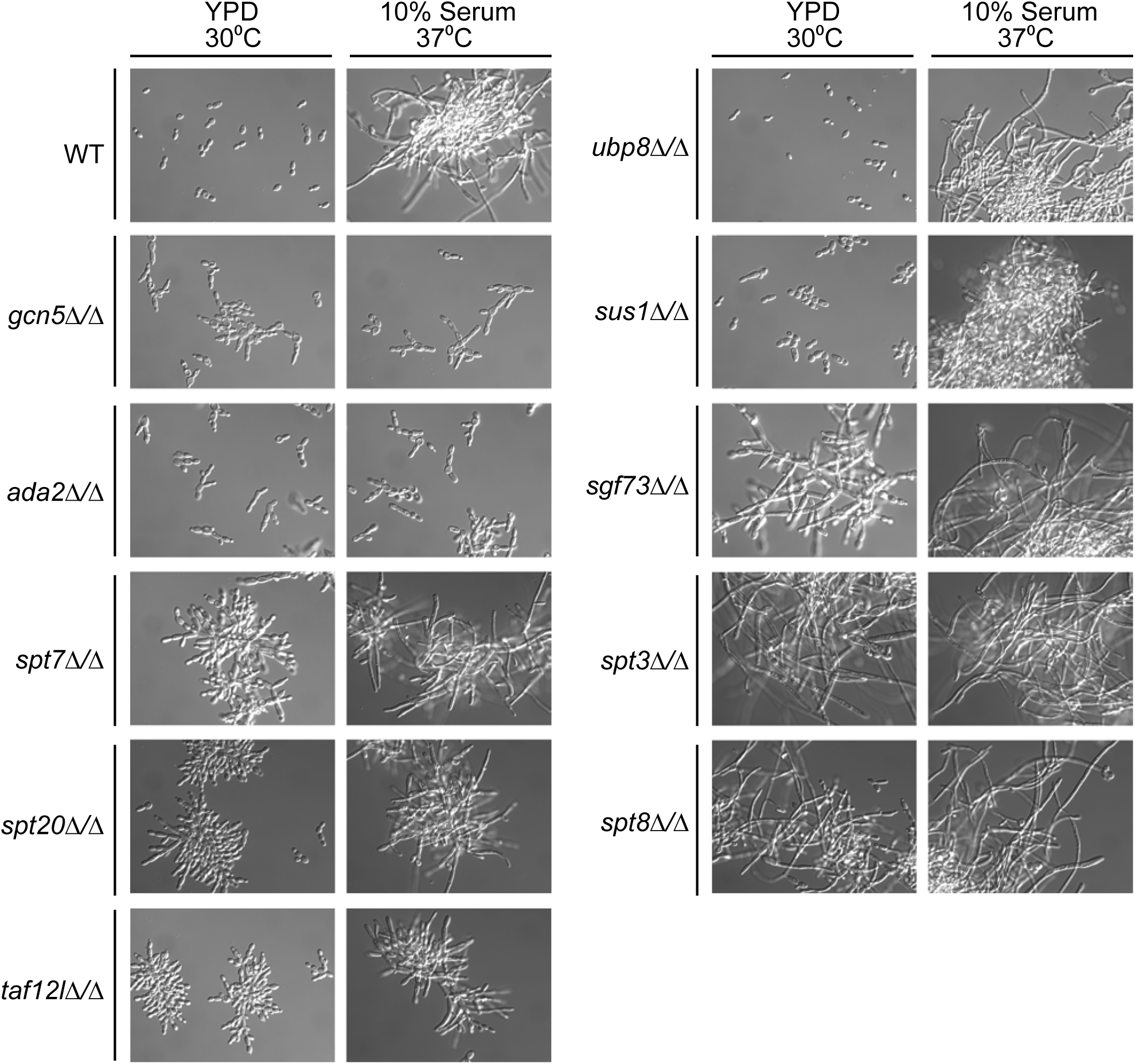
SAGA complex regulates morphological transitions of *Candida albicans.* Strains were pre-cultured overnight in YPD medium to saturation, then diluted to fresh YPD and YPD + 10% FBS and grown for 4h at 30°C and 37°C, respectively to induce hyphae formation. After 4 hours, cells were harvested, fixed with 4% formaldehyde. These cells were then mounted on glass slides and imaged under DIC mode of microscope.

Interestingly, we observed highly module-specific phenotypes across our mutants (**Figure 4**). While deletion of the HAT module subunits (*gcn5Δ/Δ, ada2Δ/Δ*) and SPT module (*spt7Δ/Δ*, *spt20Δ/Δ*) resulted in pseudohyphal growth at 30°C, deletion of TBP interaction module subunits *spt3Δ/Δ* and *spt8Δ/Δ* formed filamentous hypha-like structures under the same conditions. Furthermore, most HAT module mutant cells exhibit unipolar budding with axial branching or bipolar budding without axial branching. On the other hand, SPT module mutants formed much more pronounced, dense, highly branched aggregates (**Figure 4**). The most varied phenotypes, however, were observed for the DUB module mutants with *ubp8Δ/Δ* and *sus1Δ/Δ* mutants morphologically indistinguishable from the wild-type strain. However, *sgf73Δ/Δ* exhibited unique phenotypic traits. At 30°C, *sgf73Δ/Δ* formed elongated pseudohyphal clusters, a phenotype not observed in any other mutant (**Figure 4**). All SAGA subunit deletion strains were present in yeast form upon gene complementation (**Figure S4**).

We also tested whether the mutants are locked in a particular morphological state or can undergo morphological transformation under hypha-inducing conditions (37°C with 10% fetal bovine serum [FBS]). Interestingly, while both *gcn5Δ/Δ* and *ada2Δ/Δ* (HAT mutants) retained the pseudohyphal form and failed to transition into true hyphae, *spt7Δ/Δ* and *spt20Δ/Δ* (SPT mutants) exhibited pseudohyphal cells that are much more elongated but lack true hyphal structures (**Figure 4**). Intriguingly, *sgf73Δ/Δ* cells that formed elongated pseudo-hyphae at 30°C were able to transition to hyphal form under 37°C with 10% FBS conditions (**Figure 4**).

Colony morphology analysis on YPD and mannitol-based spider media at 30°C and 37°C gave similar results. The HAT module mutants formed smooth colonies at both temperatures, with no peripheral hyphal projections on spider medium, indicative of no hyphal transition. (**Figure S3**). Mutants in the SPT module, *spt7Δ/Δ* and *spt20Δ/Δ* mutants, formed rough colonies without the characteristic wrinkling or peripheral hyphal projections of the WT. The *spt3Δ/Δ* and *spt8Δ/Δ* mutants, by contrast, displayed rough, wrinkled colonies with visible hyphal projections at 30°C, although these projections were absent at 37°C (**Figure S3**). Colony morphology of DUB module mutants revealed that *ubp8Δ/Δ* colonies were comparable to wild-type colonies on both YPD and spider media, with much more pronounced hyphal projection. A recent study (21) reported similar phenotypes for *spt7Δ/Δ, spt8Δ/Δ* and *ubp8Δ/Δ.* However, another study on *ubp8Δ/Δ* (28) reported smooth colonies without wrinkling on YPD plates with 10% FBS. In contrast to *ubp8Δ/Δ*, *sus1Δ/Δ* did not exhibit wrinkling or filamentation on spider medium. However, *sgf73Δ/Δ* mutants consistently formed rough, wrinkled colonies across both YPD and spider media, with temperature-dependent differences. Colonies were wrinkled at both 30°C and 37°C on YPD, though hyphal projections were only observed at 37°C. On spider medium, *sgf73Δ/Δ* colonies were more wrinkled at 37°C compared to 30°C (**Figure S3**). These findings suggest that the SAGA complex regulates fungal morphogenesis through module-specific mechanisms that affect both cellular architecture and colony development.

### Dual role for SAGA complex in the transcriptional response of hypha-specific genes

The filamentation program of *C. albicans* is governed by specific transcriptional changes with upregulation of some key transcription factors such as Efg1, Brg1 and Ume6 and downstream hypha-specific genes (HSGs) such as *HWP1* and *HGC1*. So, to uncover if the SAGA subunit mutant-specific phenotypes are results of altered hyphal transcriptional gene regulation, we measured the mRNA levels of *HWP1* and *HGC1,* as well as transcriptional regulators *BRG1*, *UME6*, and *NRG1*, in various SAGA deletion mutants using qRT-PCR (**Figure 5A-B**).

**Fig. 5.**
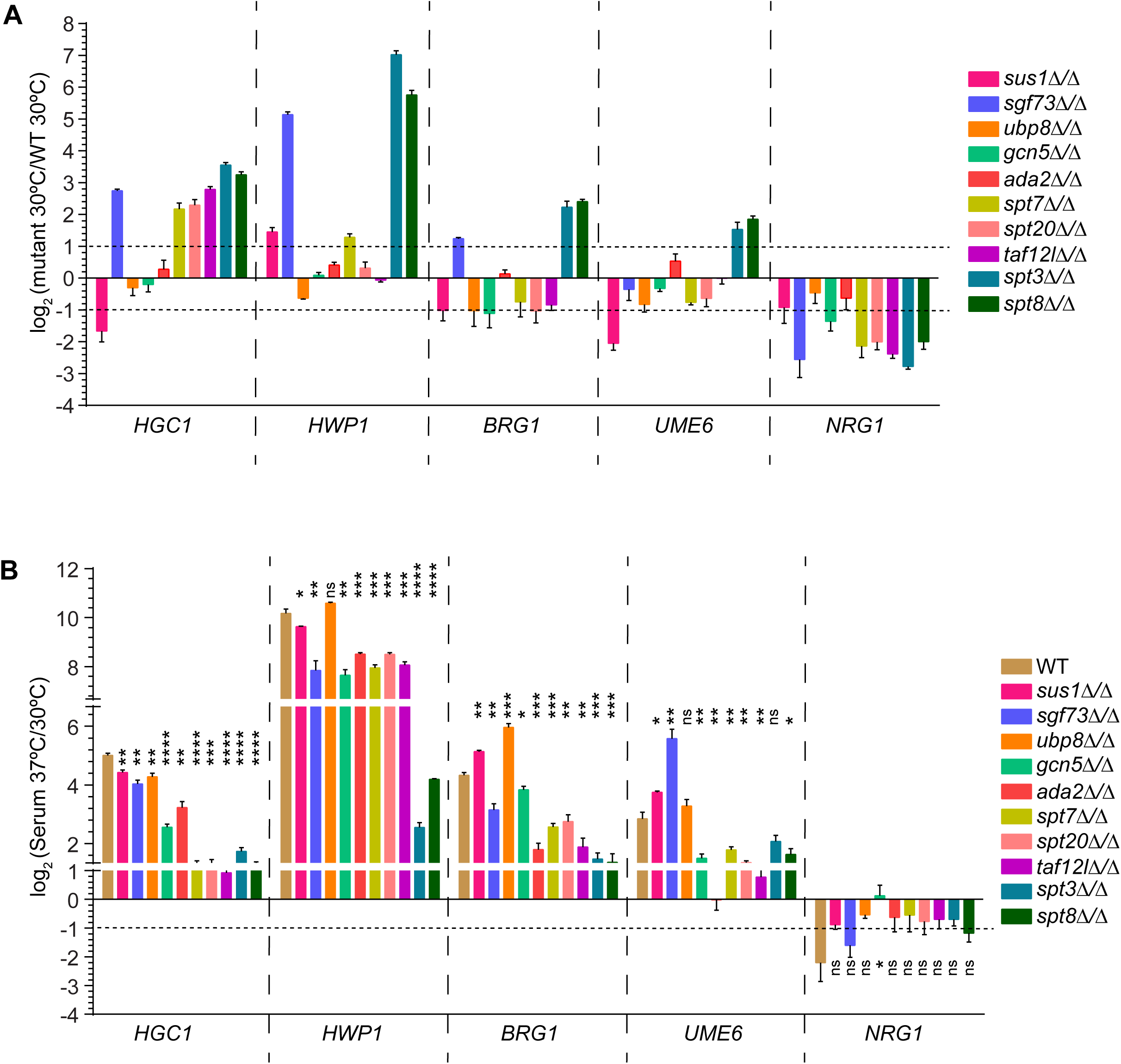
SAGA complex acts as a dual regulator of the expression of Hypha-specific genes. Quantitative RT-PCR was used to measure mRNA levels of canonical hyphal genes in WT and mutant strains under non-inducing (YPD, 30°C) and inducing (YPD + 10% FBS, 37°C) conditions. Cells were cultured similarly as described for figure 4. (A) Relative mRNA abundance in SAGA deletion mutants compared to WT in YPD at 30°C and (B) Relative mRNA abundance in WT and SAGA deletion mutants under hypha-inducing conditions (YPD with 10% FBS at 37°C) compared to non-hyphae inducing conditions (YPD, 30°C). SCR1 was used as a normalization control. Data are averages of three biological replicates. Statistical significance was determined by unpaired two-tailed t-test, with thresholds: *P* ≤ 0.05 (*), ≤ 0.01 (**), ≤ 0.001 (***), ≤ 0.0001 (****), and not significant (ns).

In WT cells at 30°C, *HWP1*, *HGC1*, *BRG1*, and *UME6* are minimally expressed, consistent with yeast-phase morphology. However, deletion of *SPT3*, *SPT8*, or *SGF73* led to strong upregulation of both *HWP1* (≥30-fold) and *HGC1* (≥6-fold) under these non-inducing conditions, consistent with the observed constitutive filamentous phenotype of these mutants. Likewise, *BRG1* and *UME6* were significantly upregulated (≥4-fold and ≥3-fold, respectively) in *spt3Δ/Δ* and *spt8Δ/Δ* strains, suggesting derepression of the upstream hyphal transcriptional program (**Figure 5A**). In the *sgf73Δ/Δ* mutant, *BRG1* showed modest upregulation (∼2-fold), but *UME6* was not significantly induced. Moreover, consistent with the morphology, *ubp8Δ/Δ* had no significant impact on hyphal gene expression, whereas *sus1Δ/Δ* caused a decrease in *HGC1* and an increase in *HWP1* (**Figure 5A**). These results support functional divergence among DUB subunits, with *SGF73* exerting a unique regulatory influence on the filamentation program. Notably, deletion of *SPT7*, *SPT20*, or *TAF12L*, structural components of the core module, led to selective upregulation of *HGC1* (≥5-fold) but not *HWP1*, indicating selective activation of the filamentation pathway. This is again consistent with the pseudo-hyphal morphology of these mutants. In contrast, deletion of *GCN5* or *ADA2*, core HAT module subunits, had no effect on *HWP1* or *HGC1* expression at 30°C (**Figure 5A**).

Next, we measured the mRNA level of these genes under filament-inducing conditions (37°C + 10% FBS). Consistent with the morphological transition of WT cells and previous studies, we observed a robust induction of *HWP1*, *HGC1*, *BRG1*, and *UME6*, whereas *NRG1* (**Figure 5B**), a known negative regulator of hyphal growth, was repressed (41). In contrast, mutants lacking *GCN5*, *ADA2*, *SPT7*, *SPT20*, *TAF12L*, *SPT3*, or *SPT8* displayed a marked decrease (≥3-fold) in the induction of *HWP1*, *HGC1*, *BRG1*, and *UME6* under these conditions (**Figure 5B**), consistent with their filamentation defects in serum at 37°C. Surprisingly, *sus1Δ/Δ* and *ubp8Δ/Δ* mutants showed increased expression of *BRG1* and *UME6* in response to serum induction. Meanwhile, *sgf73Δ/Δ* exhibited a split regulatory phenotype: decreased *BRG1* and increased *UME6* expression (**Figure 5A**). Notably, while *NRG1* levels were reduced by most SAGA mutants under non-inducing conditions, upon hyphal induction, *NRG1* levels were unaffected in all SAGA mutants (**Figure 5 A-B**), indicating that SAGA influences filamentation primarily through modulation of the *BRG1–UME6* axis rather than by repressing *NRG1*.

Together, these results reveal that the SAGA complex exerts both positive and negative regulatory effects on hypha-specific gene expression and transcription factor activation. Certain subunits (*SPT3*, *SPT8*, *SGF73*) repress filamentation gene expression under yeast conditions, while others (*GCN5*, *ADA2*, *SPT7*, *SPT20*, *TAF12L*) are required for full induction of the hyphal transcriptional program under filament-inducing conditions. These findings demonstrate that SAGA functions as a modular regulator of *C. albicans* morphogenesis, orchestrating transcriptional plasticity in response to environmental cues.

### Differential SAGA subunit contribution to Histone H3K9 acetylation

The SAGA complex exerts its transcriptional regulatory functions through two major enzymatic activities: histone acetylation, mediated primarily by the histone acetyltransferase (HAT) subunit Gcn5 (42), and histone deubiquitination, catalyzed by Ubp8 (28). To evaluate the contribution of individual SAGA subunits to histone H3 acetylation in *Candida albicans*, we assessed global H3K9 acetylation (H3K9ac) levels in a panel of SAGA subunit deletion mutants. As expected, deletion of core HAT module components *GCN5* and *ADA2* caused the most pronounced reduction in H3K9 global acetylation, with ∼85% decrease compared to the wild-type strain, confirming their essential roles in acetyltransferase activity (**Figure 6**). Deletion of the architecture or TAF module subunits, such as *TAF12L*, *SPT7*, and *SPT20,* resulted in a moderate decrease in H3K9ac levels (∼25–40%), supporting their role in maintaining SAGA structural integrity and thereby promoting enzymatic function.

**Fig. 6.**
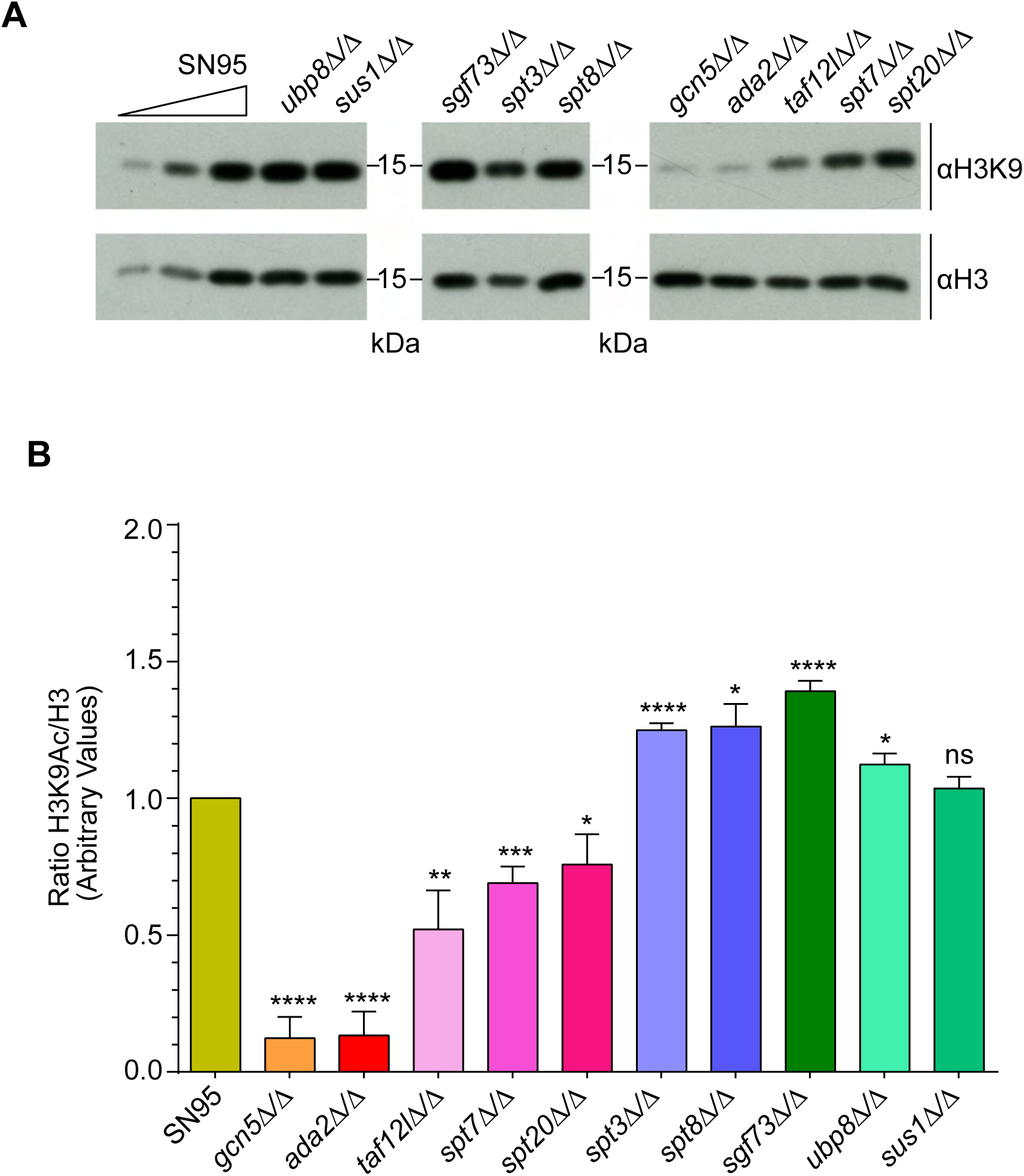
SAGA-mediated histone H3K9 acetylation depends on inter-subunit integrity. (A) Global H3K9 acetylation analysis in WT and SAGA deletion strains using western blotting. Anti-H3K9Ac and anti-H3 antibodies were used to detect acetylated and total histone H3, respectively. (B) Quantitation of bulk levels of H3K9Ac and H3 in different SAGA deletion mutants. Band intensities of Western blots were quantified from at least four replicates using ImageJ and plotted as the ratio of H3K9Ac to H3. Statistical significance was determined by an unpaired two-tailed t-test, with thresholds: *P* ≤ 0.05 (*), ≤ 0.01 (**), ≤ 0.001 (***), ≤ 0.0001 (****), and not significant (ns).

**Fig. 7.**
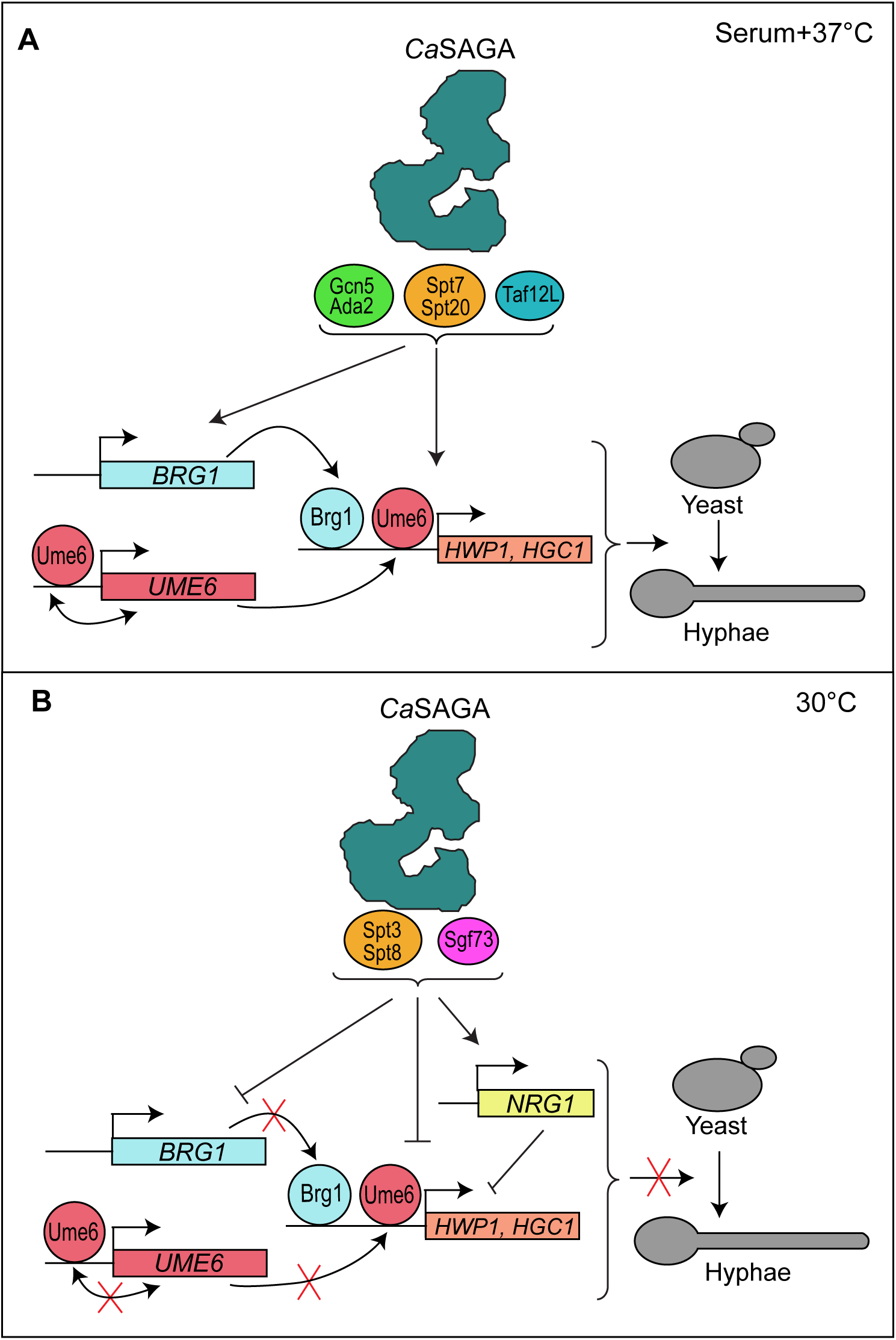
Proposed model for module-specific regulation of morphogenesis by the *Ca*SAGA complex. **(A)** Under filament-inducing conditions (e.g., serum, Spider medium, 37°C), the *Ca*SAGA complex promotes hyphal development through the coordinated action of the HAT and core scaffold modules. The HAT module (Gcn5–Ada2), together with core subunits Spt7, Spt20, and Taf12L, facilitates transcriptional activation of key hyphal regulators *BRG1* and *UME6*. Brg1 and Ume6 cooperatively induce hypha-specific effector genes such as *HWP1* and *HGC1*, driving the yeast-to-hypha transition. Under these conditions, SAGA-mediated histone acetylation supports productive transcriptional reprogramming required for sustained filamentous growth. **(B)** Under yeast-maintaining conditions (e.g., YPD at 30°C), the TBP-interaction module (Spt3–Spt8), together with the DUB-module subunit Sgf73, functions to restrain inappropriate activation of the hyphal program. This regulatory state maintains expression of the hyphal repressor *NRG1*, thereby preventing activation of *BRG1*–*UME6*–dependent transcription and preserving yeast-form growth. Loss of Spt3, Spt8, or Sgf73 disrupts this restraint, leading to derepression of hypha-specific genes and constitutive filamentation even under non-inducing conditions.

Furthermore, deletion of the deubiquitination module subunits *UBP8* and *SUS1* had little to no effect on global H3K9 acetylation, consistent with their distinct activities within different modules. Strikingly, deletion of *SGF73*, another DUB module subunit, as well as *SPT3* and *SPT8,* subunits of the TBP-interaction module, led to a surprising ∼25% increase in global H3K9ac. These findings suggest a repressive or regulatory role for these subunits in modulating the HAT activity of the SAGA complex. Collectively, these data indicate that while the enzymatic core of the HAT module is indispensable for H3K9 acetylation, other subunits, particularly those in the structural and DUB modules, play modulatory roles.

## Discussion

The evolutionarily conserved SAGA complex regulates gene expression across eukaryotes, from yeast to humans (1,2). Structural and biochemical studies have defined its modular organization comprising the Core, HAT, TBP-interaction, and DUB modules (3,5–7,43,44). Expression analyses show that subunits within the same module often affect transcription similarly (11,12,35). However, studies in filamentous fungi (e.g., *Aspergillus nidulans*, *Fusarium fujikuroi, Cryptococcus neoformans*) and plants suggest partial conservation, particularly of the DUB module (32–34,45,46).

While previous research in *Candida albicans* has explored the role of individual or subsets of SAGA subunits in gene expression and virulence (21–28) a comprehensive, module-focused analysis has been lacking. Here, we present a systematic dissection of the SAGA complex in *C. albicans*, revealing its critical role in regulating environmental stress responses, drug resistance, and morphogenesis, key factors in fungal pathogenesis. Our findings show that SAGA subunits act both independently and cooperatively to modulate their acetyltransferase and coactivator functions.

### Distinct modular activities of the SAGA complex mediate stress adaptation and antifungal resistance in *C. albicans*

Our study shows that *Candida albicans* encodes clear single orthologs for all canonical *S. cerevisiae* SAGA subunits, except *TAF12* (26). A previous study (22) listed SAGA subunits in *C. albicans;* however, unlike our study, it did not identify *SGF73* (ORF 19.3473). We later reported two *TAF12* paralogs, including *TAF12L* (ORF 19.470), as part of SAGA (26). Domain analysis confirms structural conservation across key subunits and modules HAT, DUB, TBP-interaction, and Core, indicating that *C. albicans* harbors a complete and evolutionarily conserved SAGA complex.

Of the eleven SAGA subunits tested, nine (*GCN5, ADA2, UBP8, SUS1, SGF73, SPT7, SPT20, TAF12L, SPT3, and SPT8*) were nonessential for viability, while *TRA1* and *ADA1* seems to be essential. *TRA1* ATP-binding cleft mutant was found to regulate drug resistance and filamentation in *C. albicans* (47). However, deletion of all except *UBP8* resulted in significant growth defects under nutrient-rich conditions, underscoring the roles of these genes in cellular homeostasis. Phenotypic profiling across a range of environmental stress conditions revealed distinct, module-specific roles for SAGA in stress adaptation. Deletion mutants of the HAT were consistently hypersensitive to oxidative stress, iron limitation, amino acid deprivation, and heavy metal exposure. Similar phenotypes were observed in core module (*SPT* subunits and *TAF12L*) mutants, consistent with the core module’s architectural and scaffolding roles, which are critical for complex stability and activity (1). Our comparative analysis of homologues *in S. cerevisiae* SAGA mutants revealed very similar phenotypic profiles, indicating functional conservation of inherent activities across the two fungal species. Notably, the DUB module showed the most intra-modular variation. The *sgf73Δ/Δ* mutant in *C. albicans* exhibited severe stress-related defects, unlike the *ubp8Δ/Δ* or *sus1Δ/Δ* mutants. In contrast, *S. cerevisiae sgf73Δ* displayed phenotypes similar to other DUB subunit deletions. This suggests a distinct role for *CaSGF73* beyond its known DUB function, consistent with studies in *S. pombe* and humans implicating *SGF73* in additional pathways (48–51). Interestingly, fluconazole sensitivity appears to be module-dependent: *SPT* deletions increased susceptibility, whereas HAT mutants (*gcn5Δ/Δ*, *ada2Δ/Δ*) showed resistance, in contrast to earlier findings (21,22) This discrepancy may reflect strain or media differences and highlights the complex, context-dependent role of SAGA in drug response. Altogether, our findings position the *C. albicans* SAGA complex as a modular and dynamic coactivator that integrates environmental signals to modulate transcriptional responses critical for fungal stress tolerance and virulence.

### Modulation of morphogenetic transitions by the SAGA complex in *C. albicans*

The ability of *Candida albicans* to switch between yeast, pseudohyphal, and hyphal forms is central to its pathogenicity, facilitating both systemic and invasive infections (39,52,53). While much is known about transcriptional regulators of morphogenesis, chromatin-modifying mechanisms, particularly histone acetylation, are emerging as key modulators of these transitions (54–57). Interestingly, our data reveal that the SAGA complex plays both activating and repressive roles in *C. albicans* morphogenesis. Deletion of HAT module subunits (*gcn5Δ/Δ*, *ada2Δ/Δ*) and core components (*spt7Δ/Δ*, *spt20Δ/Δ*, *taf12lΔ/Δ*) resulted in pseudohyphal morphology at 30°C and impaired hyphal induction in serum and Spider media. Conversely, deletion of TBP-interacting subunits (*spt3Δ/Δ*, *spt8Δ/Δ*) led to constitutive filamentation, even under non-inducing conditions, suggesting a repressive role for this module in maintaining yeast form. A similar dual role for SAGA has been reported in *S. pombe,* where Gcn5 and Spt8 function in opposing pathways that regulate sexual differentiation (31). Notably, *sgf73Δ/Δ* exhibited an elongated pseudohyphal phenotype distinct from that of *sus1Δ/Δ* and *ubp8Δ/Δ,* yet it still responded to serum-induced hyphal cues, suggesting a non-canonical role. Transcriptional analysis confirmed dysregulation of hypha-specific effectors (*HWP1*, *HGC1*) and key upstream regulators (*BRG1*, *UME6*) in these mutants, while *NRG1* expression remained largely unaffected. This suggests that SAGA primarily modulates morphogenesis through *BRG1* and *UME6*, rather than by repressing *NRG1*.

### DUB-module subunit *SGF73* plays a non-canonical unique role in *C. albicans*

SGF73 (the yeast ortholog of human ATXN7) is a crucial subunit of the SAGA complex in *Saccharomyces cerevisiae*, playing an essential role in the activation of the deubiquitination (DUB) module. Structural studies of the SAGA complex (3,5,44,58) have demonstrated that Sgf73 anchors the DUB module to the core SAGA complex and facilitates the coordination between its assembly and catalytic activity. Interestingly, *SGF73* is absent from plants (46) and has not been identified in *Cryptococcus neoformans* (32), along with other DUB module components. In yeast and other organisms, *SGF73* has also been reported to function independently of the SAGA complex and H2B deubiquitination (59). Our findings show that the *Candida albicans sgf73Δ/Δ* null mutant displays pronounced stress sensitivity, resembling phenotypes observed in core module subunit deletion mutants. Additionally, the *sgf73Δ/Δ* mutant exhibits a hyper-filamentous phenotype at 30°C, closely resembling that of TBP-interaction module mutants such as *spt3Δ/Δ* and *spt8Δ/Δ*. Notably, the *sgf73Δ/Δ* mutant also shows elevated global levels of H3K9 acetylation, indicating that loss of *SGF73* may indirectly affect histone acetylation dynamics (discussed below). Gcn5 is known to acetylate the Sgf73 subunit at different lysine residues, suggesting post-translational modification-based interaction between the HAT and DUB module subunits (60). Collectively, these results suggest that *CaSGF73* may have a non-canonical role in *C. albicans*, potentially independent of the SAGA DUB module. Further investigation is needed to clarify the nature and significance of this role.

### Interplay between SAGA complex modules regulate H3K9 acetylation in *C. albicans*

SAGA-mediated histone acetylation plays a pivotal role in regulating stress responses and morphogenesis in *Candida albicans* (24,27,61). In *S. cerevisiae*, while Gcn5 functions as the catalytic subunit, full acetyltransferase activity requires additional HAT and SPT module components. To explore this in *C. albicans*, we assessed global histone H3 lysine-9 acetylation (H3K9ac) across various SAGA subunit deletion mutants. As expected, HAT module mutants (*gcn5Δ/Δ*, *ada2Δ/Δ*) showed a marked reduction in H3K9ac, confirming their essential role in acetylation. Core module mutants (*spt7Δ/Δ*, *spt20Δ/Δ*, *taf12lΔ/Δ*) exhibited moderate decreases, suggesting conserved structural contributions to HAT function. Interestingly, deletion of TBP-interacting subunits (*spt3Δ/Δ*, *spt8Δ/Δ*) led to increased H3K9ac levels, suggesting a previously unrecognized role in restricting hyperacetylation, although the underlying mechanism remains unclear. A similar hyperacetylation phenotype was observed in the *sgf73Δ/Δ* mutant, aligning with *S. cerevisiae* studies showing that *SGF73* mutations can variably affect SAGA’s HAT activity (58). Given structural evidence that the HAT and DUB modules are closely positioned (3,5), these findings suggest intermodular regulation of SAGA’s enzymatic activity.

Overall, our study highlights the complex interplay between SAGA modules in modulating histone acetylation and underscores their broader roles in *C. albicans* growth, morphogenesis, and stress adaptation.

## Conclusion

In this study, we comprehensively explored the functional roles of the SAGA complex in *Candida albicans*, revealing its critical involvement in regulating multiple aspects of fungal biology, including morphogenesis, stress response, and antifungal resistance. Our systematic deletion analysis of various SAGA subunits demonstrated that while some subunits are non-essential for viability, they are indispensable for efficient growth under stress conditions. Moreover, our findings underscore the modular nature of the SAGA complex, with distinct modules (HAT, SPT, DUB) playing specialized roles in regulating gene expression and acetylation, particularly during the *C. albicans* transition from yeast to hyphal forms, which is crucial for pathogenicity. Notably, we identified significant interactions among subunits, elucidating how these interactions regulate critical transcription factors and histone marks, such as H3K9 acetylation, that govern the expression of genes involved in morphogenesis and stress adaptation. These insights into the molecular mechanisms governing SAGA function in *C. albicans* contribute to a broader understanding of fungal pathogenesis.

## Materials and Methods

### Strains, plasmids, and oligonucleotides

Plasmids, strains, and oligonucleotides used in this study are listed in Supplemental Tables S2, S3, and S4.

### Media and growth conditions

*C. albicans* and *S. cerevisiae* strains were routinely cultured in yeast extract-peptone-dextrose (YPD) medium (1% yeast extract, 2% peptone, 2% dextrose) at 30°C unless otherwise specified. To study stress resistance and morphogenesis, strains were plated on various media, including SPIDER medium (0.1% yeast extract, 0.1% peptone, 0.1% glucose, 2% agar) and synthetic complete (SC) medium. Stress assays were performed by supplementing YPD with stress-inducing agents as indicated in Table S5.

### Identification of *C. albicans* SAGA subunits

SAGA subunit orthologs in *C. albicans* were identified by sequence similarity to their counterparts in *Saccharomyces cerevisiae* using the Basic Local Alignment Search Tool (BLAST). Domain prediction for each subunit was performed using the SMART tool (30) with further validation through sequence alignment with *S. cerevisiae* homologs. For the *CaSpt3* and *CaAda1* subunits, predictions were cross validated with the Phyre2 tool (62) to assign specific domain coordinates. For the *CaSgf73* subunit, the Zinc Finger domain (amino acids 55-109) was predicted using Phyre2, as the SMART tool did not detect it. This domain prediction was further validated through sequence alignment with *S. cerevisiae* Sgf73, confirming the domain’s presence in *C. albicans*.

### Construction of *C. albicans* SAGA deletion mutant strains

Gene-specific disruption cassettes were generated using the *pHAH1* plasmid (63)a modified version of the single-transformation gene deletion cassette (64). The deletion strategy employed a split-marker technique (65), in which primers corresponding to the regions flanking the target gene were used to amplify up- and down-split fragments with approximately 1.0-kb overlaps within the *ARG4* gene. The resulting cassette was transformed into *C. albicans* strain SN95, and Arg^+^ transformants were selected. After transformation, PCR was used to confirm the correct integration of the deletion cassette. Second-allele replacements were selected based on growth on SC plates lacking His and Arg. Homozygous deletion mutants were then confirmed by PCR amplification of the targeted loci using locus-specific primers and ORF-specific primers to exclude the presence of any additional copies of the target genes.

### Complementation of *C. albicans* SAGA deletion mutant strains

For reintegration of *GCN5*, *SPT7*, *SPT20, SUS1, SGF73, ADA2, SPT3 and SPT8* into the homozygous deletion strains SDC57, SDC67, SDC69, SDC63, SDC65, SDC59, SDC73 and SDC71, plasmid pNIM1 was sequentially digested with BglII/MluI to obtain nourseothricin resistance marker CaSAT1, while plasmid pCip10 was digested with NotI/XbaI and end filled with Klenow. The linearized plasmid pCip10 and insert CaSAT1 was ligated and transformed into electrocompetent bacterial cells DH10B to obtain plasmid pCip10-SAT1 (pPI1). Then, genes were amplified along with 1000bp upstream (1500bp upstream for *SPT7 ORF)* and 500bp downstream region of ORF sequences using primer pairs ONC1156-ONC1157, ONC1152-ONC1153, ONC1154-ONC1155, ONC1158-ONC1159, ONC1160-ONC1161, ONC1162-ONC1163, ONC1164-ONC1165, ONC1166-ONC1167 for *GCN5*, *SPT7, SPT20, SUS1, SGF73, ADA2, SPT3* and *SPT8* respectively. The amplified product was digested with BamHI (for *SPT7*, *SPT20, SUS1, SGF73, ADA2, SPT3* and *SPT8*) and SacI (for *GCN5*) alongwith plasmid pPI1 to obtain plasmid pPI2 (*GCN5*), pPI3 (*SPT7*), pPI4 (*SPT20*), pPI5 (*SUS1*), pPI6 (*SGF73*), pPI7 (*ADA2*), pPI8 (*SPT3*) and pPI9 (*SPT8*) respectively.

Then obtained plasmids pPI2, pPI3, pPI4, pPI5, pPI6, pPI7 and pPI8 were linearized with PmeI, BstEII, AflII, MscI, PmlI, BmgBI, and NsiI respectively, and integrated at their native loci in mutant strains SDC57, SDC67, SDC69, SDC63, SDC65, SDC59, SDC73 to obtain complemented strains PRI1 (*gcn5Δ/Δ::GCN5*), PRI2 (*spt7Δ*/*Δ::SPT7*), PRI3 (*spt20Δ/Δ::SPT20*), PRI4 (*sus1Δ*/*Δ::SUS1*), PRI5 (*sgf73Δ/Δ::SGF73*), PRI6 (*ada2Δ/Δ::ADA2*), PRI7 (*spt3Δ*/*Δ::SPT3*). The plasmid pPI9 was digested with BglII and integrated at the RP10 locus in the mutant SDC71 strain to obtain the complemented strain PRI8 (*spt8Δ/Δ::SPT8*). The transformants were screened by PCR using ORF-specific primer pairs ONC661/ONC662 (*GCN5)*, ONC663/ONC664 (*SPT7)*, ONC708/ONC709 (*SPT20)*, ONC730/ONC731 (*SUS1)*, ONC734/ONC735 (*SGF73)*, ONC736/ONC737 (*ADA2)*, ONC732/ONC733 (*SPT3)* and ONC794/ONC795 (*SPT8)*, and the correct integration confirmed.

### RNA extraction and qRT-PCR analysis

*Candida albicans* strains were grown overnight in YPD medium and then diluted into either fresh YPD medium or YPD supplemented with 10% fetal bovine serum (FBS) for 4 hours at 30°C or 37°C, conditions that induce different forms of stress and morphogenetic transitions. Total RNA was extracted using the hot phenol method as previously described (66) as described previously (67), and the concentration was determined using a NanoDrop spectrophotometer. Total RNA was treated with RNase-free DNase I (Invitrogen) to remove genomic DNA contamination and reverse-transcribed into cDNA using the high-capacity cDNA reverse transcription kit (Invitrogen). Quantitative RT-PCR was performed using SYBR Green PCR Master Mix (Applied Biosystems) on an Applied Biosystems 7500 real-time PCR system. Relative gene expression levels were calculated using the comparative cycle threshold (CT) method (2^ΔΔCT^) (68) with *SCR1* RNA serving as an endogenous control for normalization. Control reactions were carried out in the absence of reverse transcriptase to confirm the absence of DNA contamination.

### Preparation of *C. albicans* cells for microscopy

To assess morphogenetic transitions, strains were grown in YPD or YPD+10% FBS at 30°C or 37°C, which induces yeast-to-hypha transition at the higher temperature. Cells were fixed with 4% formaldehyde (prepared in 1X PBS) for 30 minutes at room temperature (RT) and then washed five times with 1X PBS to remove residual formaldehyde. The fixed cells were mounted on glass slides and imaged using a Nikon Eclipse 90i microscope with a 60X oil immersion objective to capture morphological details. Images were processed using NIS-Elements software (Nikon).

### Colony morphology imaging

For colony morphology assessment, strains were cultured in YPD or SPIDER medium and incubated at 30°C or 37°C for the specified times (2 days for YPD, 4 days for SPIDER). Isolated colonies were imaged using a Nikon SMZ1500 stereo zoom microscope at 1X, 3X, and 5X magnification. For 5X zoom images, transmitted light was used to capture clear morphological features. Colony size, shape, and texture were used to assess the effect of gene deletions on colony formation.

### Western blotting for Histone acetylation

Histone acetylation levels were measured by Western blotting. Strains were grown overnight in YPD medium, then diluted into fresh YPD and incubated for 6 hours to mid-log phase. Cells were harvested by filtration, washed with sterile water, and flash-frozen in liquid nitrogen. The wet weight of the cell pellet was recorded, and the cells were resuspended in NI buffer (0.25 M sucrose, 60 mM KCl, 5 mM MgCl2, 1 mM CaCl2, and 0.8% Triton X-100). Cell lysis was performed by vortexing with 0.5 mm glass beads for 15 minutes at 4°C, followed by two washes of the beads with NI buffer. The lysate was clarified by centrifugation at 13,000 rpm, and the pellet was resuspended in SDS-PAGE sample buffer and boiled. Proteins were separated on a 15% SDS-PAGE gel and transferred to a nitrocellulose membrane. Membranes were probed with primary antibodies against H3K9 acetylation (anti-H3K9Ac, 07-352 Merck Millipore) or total histone H3 (anti-H3, Ab1791 Abcam). After incubation with secondary antibodies, proteins were visualized using an ECL substrate and captured on a chemiluminescence imaging system (Bio-Rad).

## Funding

Junior and Senior Research Fellowships from CSIR supported PP, PN, and MKS. K.N. acknowledges research funding from the Science and Engineering Research Board (EMR/2017/000161; CRG/2022/005145), and departmental funding support under the DBT-BUILDER (BT/INF/22/SP45382/2022) and DST-FIST II grants.

## Author Contributions

PP and KN conceptualized the study and analysed the data. PP, MKS, and PN performed the experiments. PP, MKS, and KN analyzed data and drafted the manuscript. PP, PN, MKS, and KN edited the manuscript. K.N. acquired funding, supervised the project, and finalized the manuscript.

## Acknowledgements

We thank members of the Natarajan laboratory for helpful comments and suggestions.

## Author Notes

Conflict of Interest

The author(s) declare no conflict of interest.

## Supplementary Figure Legends

**Fig. S1. Complementation of *C. albicans* SAGA mutants.** Spot assay showing growth phenotype of WT, CaSAGA deletion mutant and the complemented strains. All strains, including the WT strain SN95, were grown in YPD liquid medium till saturation, and serial dilutions were spotted on YPD, YPD + 90µM Menadione, YPD + 2.5µg/ml Fluconazole, SC-LIM, and SC-LIM + 100µM BPS plates. The plates were incubated at 30°C and images acquired.

**Fig. S2. SAGA complex is required for the stress response in *S. cerevisiae.*** (A) Spot assay showing growth phenotype of different ScSAGA deletion mutant strains. All mutants and the WT strain BY4741 were grown in YPD liquid medium until saturation, and six serial dilutions were spotted onto YPD plates. The plates were incubated at both 30°C and 37°C and imaged after every 12h. The growth phenotype shown is after 24h of growth.

**Fig. S3. Colony morphology of *C. albicans* SAGA deletion mutants.** Colony morphology of the different CaSAGA deletion mutant strains and the WT *C. albicans* strain SN95 on YPD and hyphae-inducing Spider medium. The mutants and the WT strain were grown in YPD liquid medium until saturation, and ∼200 cells were plated on the indicated plates. The plates were incubated at 30°C and 37°C, and isolated colonies were imaged on a Nikon Stereozoom SMZ1500 at 3x and 5x magnification under epi- and trans-illumination, respectively. Colonies on YPD plates were imaged after 2d and 4d growth, respectively.

---

**Table S1:** Composition of SAGA complex in different Fungal species. Identified SAGA complex subunits for different fungal species based on computational analysis and biochemical complex purifications. *Sc*SAGA: *Saccharomyces cerevisiae* SAGA (58,70)*Sp*SAGA: *Schizosaccharomyces pombe* SAGA (31), *An*SAGA: *Aspergillus nidulans* SAGA (34), *Ff*SAGA: *Fusarium fujikuori* SAGA (33), *Ca*SAGA: *Candida albicans* SAGA (22,29) and this work and *Cn*SAGA: *Cryptococcus neoformans* SAGA (32)* Identified only by computational analysis, not by biochemical purifications. ** Identified by computational analysis (36,71), N.I. – not identified. For *Candida albicans,* systematic names as well as the functionally annotated names are given.

| **SAGA modules** | **ScSAGA Subunits** | **SpSAGA Subunits** | **AnSAGA Subunits** | **FfSAGA Subunits** | **CaSAGA Subunits\*** | **CnSAGA Subunits\*** |
| --- | --- | --- | --- | --- | --- | --- |
| HAT module | *GCN5* | SPAC 1952.05 | AN3621 | FFUJ_00382 | *GCN5* orf19.705 | CNAG_00375 |
|  | *ADA2* | SPCC 24B10.08c | AN10763 | FFUJ_08807 | *ADA2* orf19.2331 | CNAG_01626 |
|  | *ADA3* | SPBC 28F2.10c | AN0440 | FFUJ_00496 | *NGG1* orf19.3023 | CNAG_04674 |
|  | *SGF29* | SPBC 1921.07c | AN0668 | FFUJ_09968 | orf19.7074 | CNAG_06392 |
| DUB module | *UBP8* | SPAC 13A11.04c | AN3711* | FFUJ_13672* | orf19.1767 | N.I. |
|  | *SGF73* | SPCC 126.04c | AN11747* | FFUJ_08521* | orf19.3473 | N.I. |
|  | *SGF11* | SPAC 57A10.14 | AN8685* | FFUJ_06279* | orf19.7360 | N.I. |
|  | *SUS1* | SPBC 6B1.12c | AN7253* | N.I. | orf19.6795 | CND_05070 |
| Activator interaction module | *TRA1* | SPBP 16F5.03c | AN8000 | FFUJ_13284 | *TRA1* orf19.139 | CNAG_07377 |
| TBP interaction module | *SPT3* | SPCC 61.02 | AN0719 | FFUJ_05029 | *SPT3* orf19.7622 | CNAG_05290 |
|  | *SPT8* | SPBC 14C8.17c | AN4670 | FFUJ_05575 | orf19.4312 | CNAG_06597 |
| Structural integrity module | *ADA1* | SPBC 887.18c | AN10953 | FFUJ_07552 | orf19.307 | CNAG_06852** |
|  | *SPT7* | SPBC 25H2.11c | AN4894 | FFUJ_13527 | *SPT7* orf19.7572 | CNAG_03850 |
|  | *SPT20* | SPAC 4D7.10c | AN0976 | FFUJ_01790 | *SPT20* orf19.422 | CNAG_04512** |
| TAF module | *TAF5* | SPCC 5E4.03c | AN0292 | FFUJ_13741 | orf19.536 | CNAG_05428 |
|  | *TAF6* | SPCC 16C4.18c | AN8232 | FFUJ_00679 | *TAF60* orf19.7454 | CNAG_02536 |
|  | *TAF9* | SPAC 12G12.05c | AN0794 | FFUJ_08952 | orf19.1111 | CNAG_07565 |
|  | *TAF10* | SPBC 21H7.02 | AN0154 | FFUJ_01846 | orf19.3242 | CNAG_01972 |
|  | *TAF12* | SPAC 15A10.02 | AN2769 | FFUJ_13236 | *TAF12L* orf19.470 | CNAG_06861 |

**Table S2:** List of Strains.

| STRAINS | RELEVANT GENOTYPE | SOURCE |
| --- | --- | --- |
| SN95 | *arg4Δ/arg4Δ his1Δ/his1Δ URA3/ura3Δ::imm<sup>434</sup>*<br>*IRO1/iro1Δ::imm<sup>434</sup>* | (65) |
| ISC36 | SN95 *taf12l Δ::FRT/ taf12lΔ::SAT1 FLP* | (26) |
| SDC56 | SN95 *gcn5Δ::HAH1/GCN5* | This work |
| SDC57 | SN95 *gcn5Δ::HAH1/gcn5Δ::HIS1* | This work |
| SDC58 | SN95 *ada2Δ::HAH1/ADA2* | This work |
| SDC59 | SN95 *ada2Δ::HAH1/ada2Δ::HIS1* | This work |
| SDC60 | SN95 *ubp8Δ::HAH1/UBP8* | This work |
| SDC61 | SN95 *ubp8Δ::HAH1/ubp8Δ::HIS1* | This work |
| SDC62 | SN95 *sus1Δ::HAH1/SUS1* | This work |
| SDC63 | SN95 *sus1Δ::HAH1/sus1Δ::HIS1* | This work |
| SDC64 | SN95 *sgf73Δ::HAH1/SGF73* | This work |
| SDC65 | SN95 *sgf73Δ::HAH1/sgf73Δ::HIS1* | This work |
| SDC66 | SN95 *spt7Δ::HAH1/SPT7* | This work |
| SDC67 | SN95 *spt7Δ::HAH1/spt7Δ::HIS1* | This work |
| SDC68 | SN95 *spt20Δ::HAH1/SPT20* | This work |
| SDC69 | SN95 *spt20Δ::HAH1/spt20Δ::HIS1* | This work |
| SDC70 | SN95 *spt8Δ::HAH1/SPT8* | This work |
| SDC71 | SN95 *spt8Δ::HAH1/spt8Δ::HIS1* | This work |
| SDC72 | SN95 *spt3Δ::HAH1/SPT3* | This work |
| SDC73 | SN95 *spt3Δ::HAH1/spt3Δ::HIS1* | This work |
| SDC74 | SN95 *ada1Δ::HAH1/ADA1* | This work |
| SDC75 | SN95 *tra1Δ::HAH1/TRA1* | This work |
| PRI1 | *gcn5Δ::HAH1/gcn5Δ::HIS1 <pPI2>* | (27) |
| PRI2 | *spt7Δ::HAH1/spt7Δ::HIS1 <pPI3>* | (27) |
| PRI3 | *spt20Δ::HAH1/spt20Δ::HIS1 <pPI4>* | (27) |
| PRI4 | *sus1Δ::HAH1/sus1Δ::HIS1 <pPI5>* | This work |
| PRI5 | *sgf73Δ::HAH1/sgf73Δ::HIS1 <pPI6>* | This work |
| PRI6 | *ada2Δ::HAH1/ada2Δ::HIS1 <pPI7>* | This work |
| PRI7 | *spt3Δ::HAH1/spt3Δ::HIS1 <pPI8>* | This work |
| PRI8 | *spt8Δ::HAH1/spt8Δ::HIS1 <pPI9>* | This work |

**Table S3:**
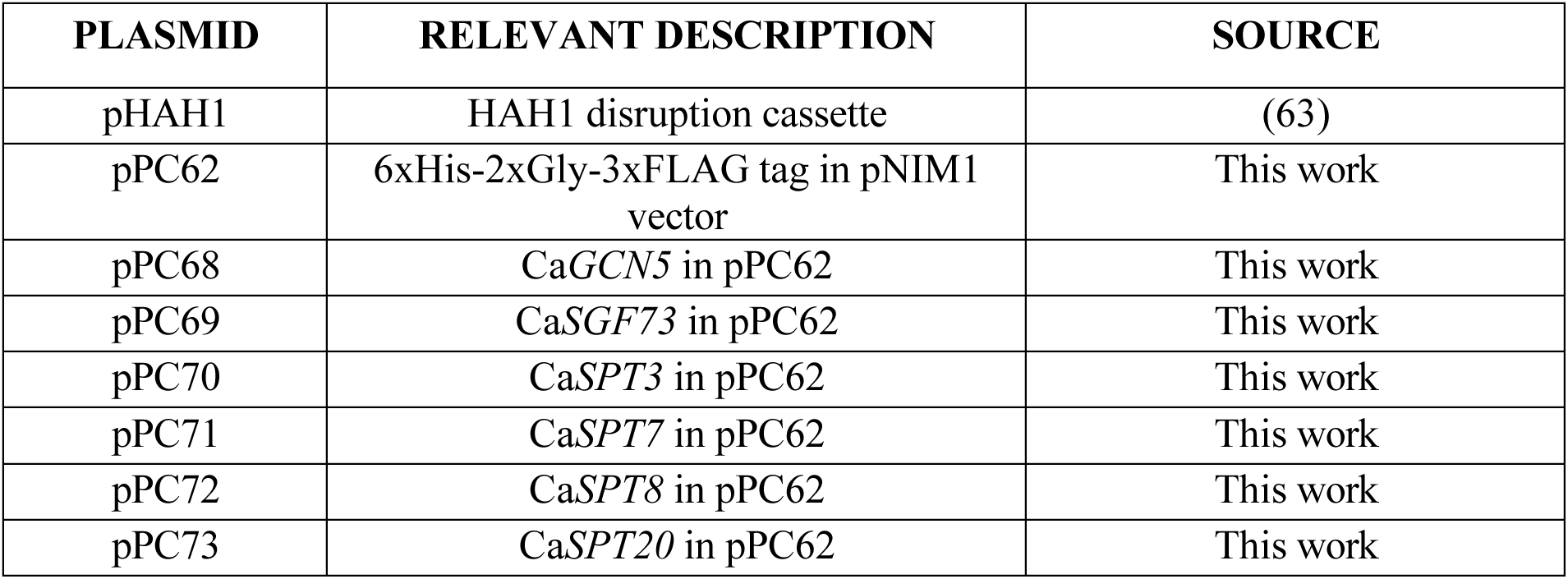
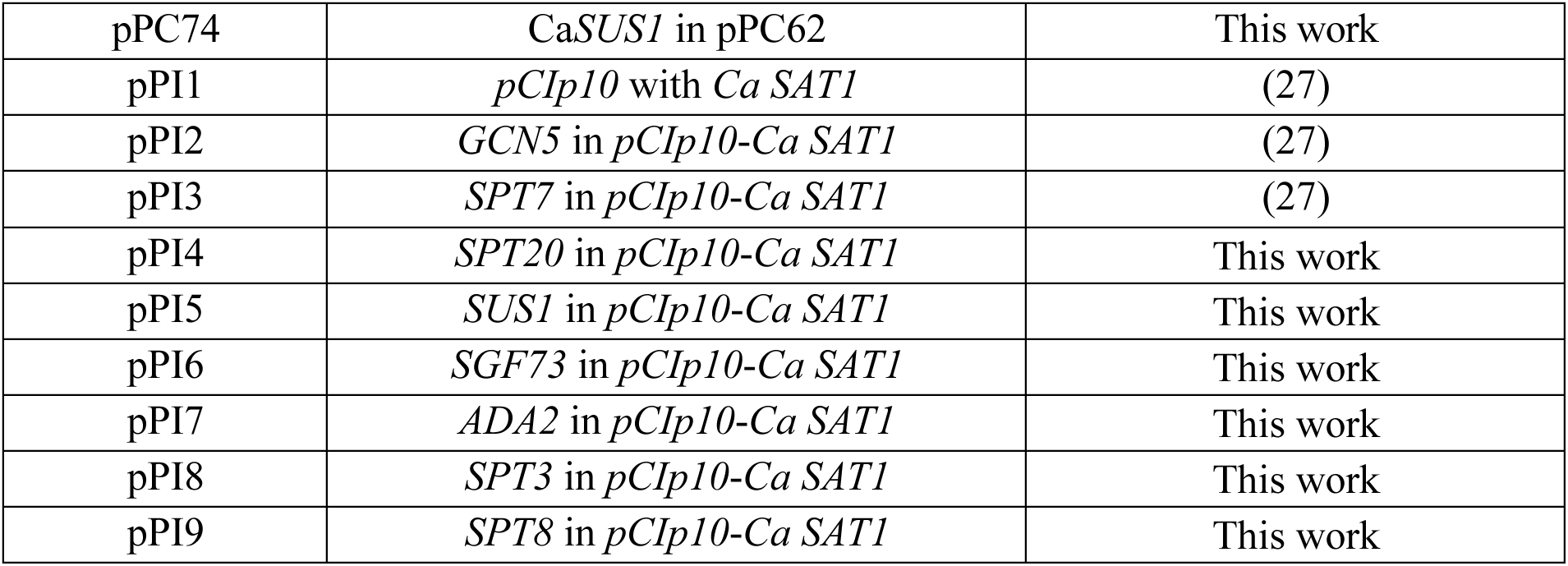
List of Plasmids.

**Table S4:**
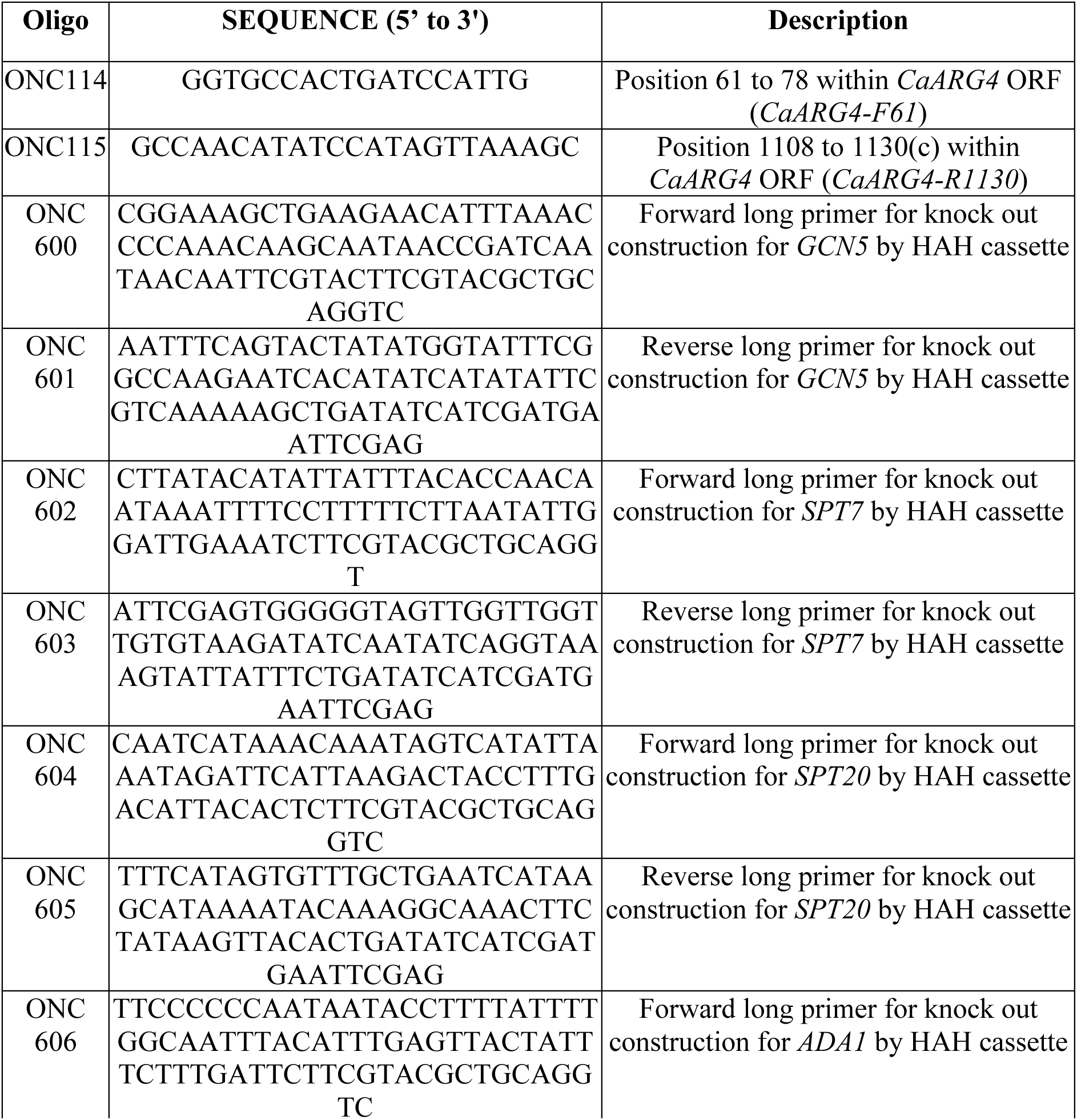
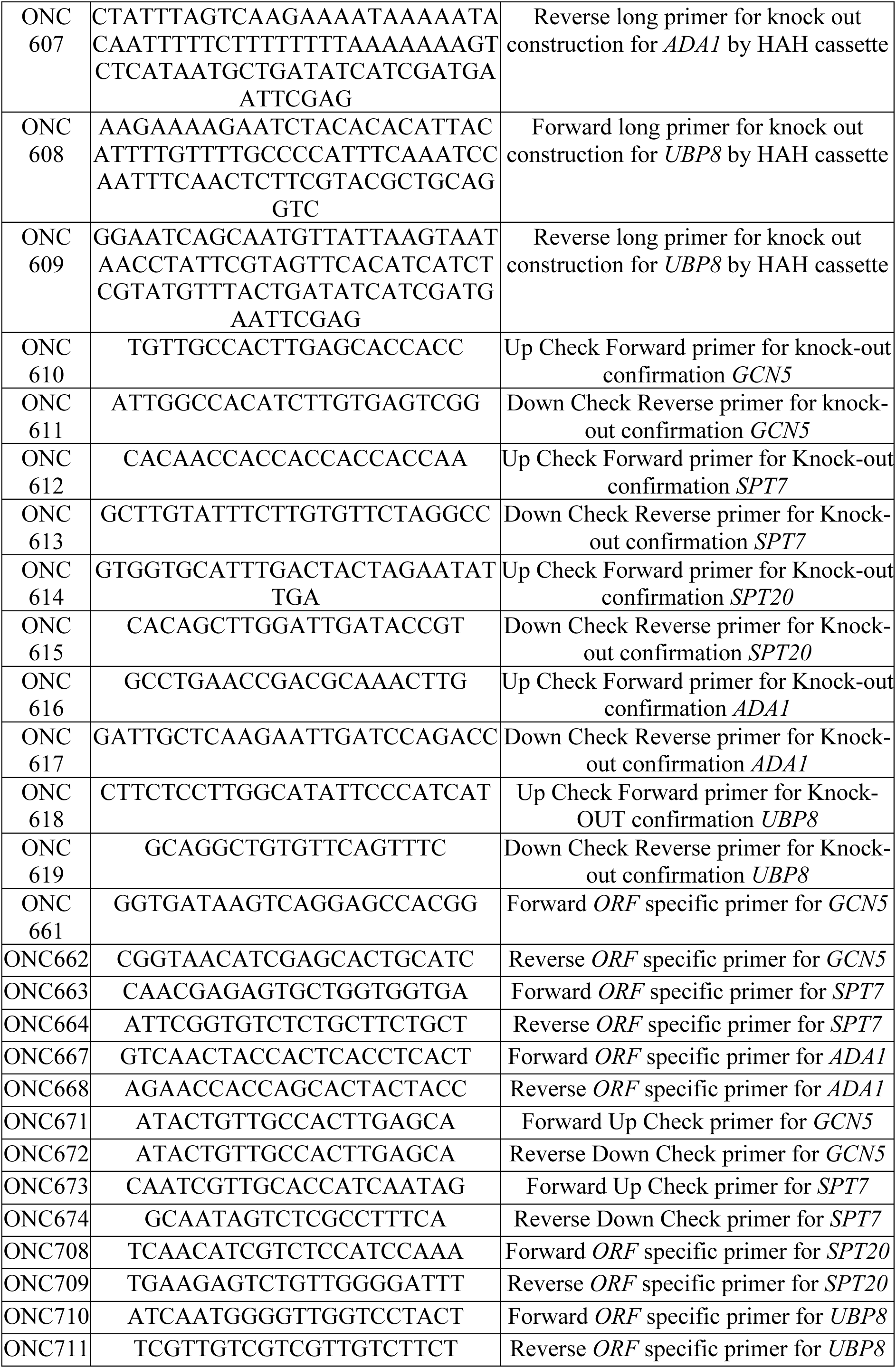
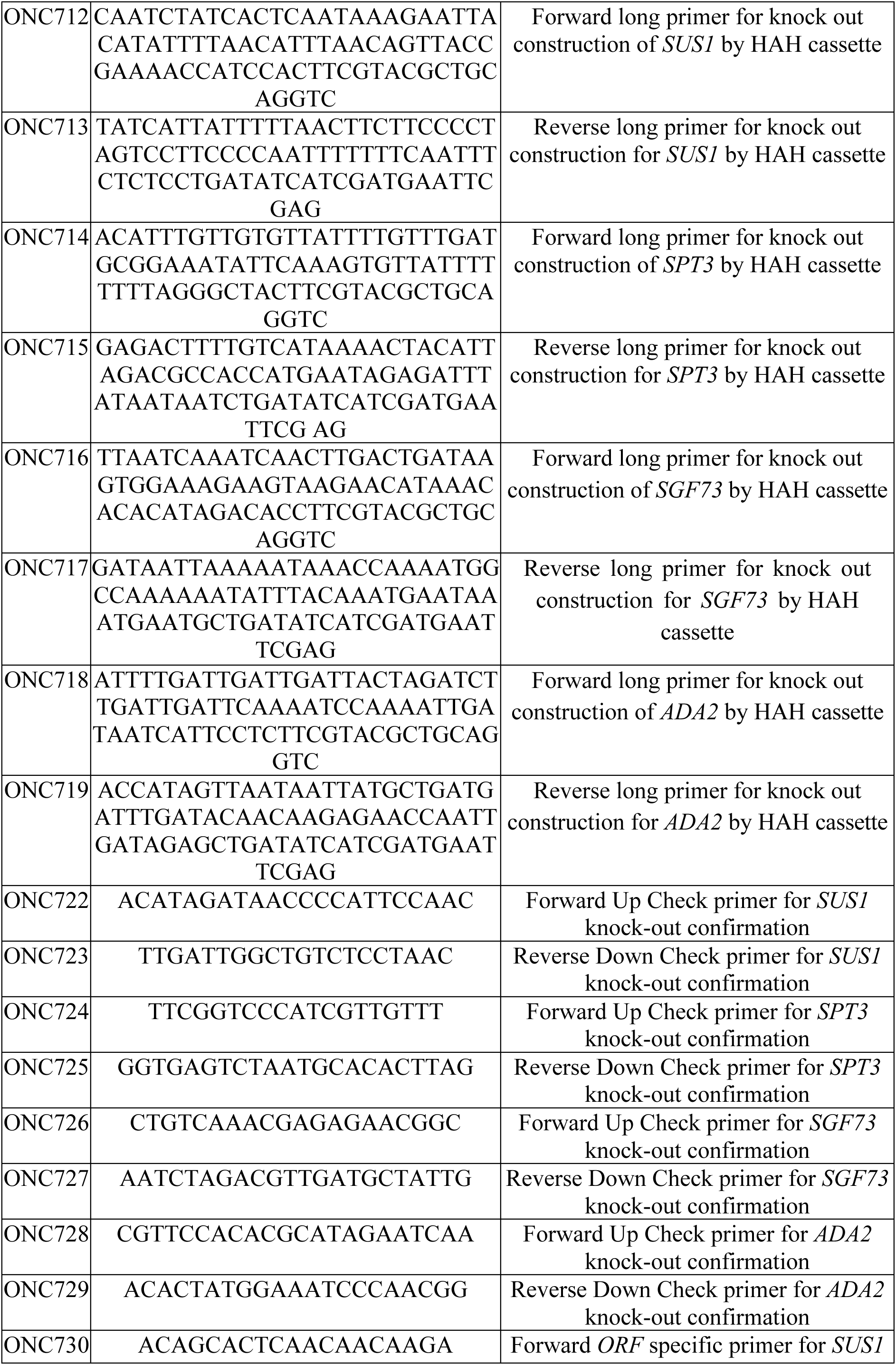
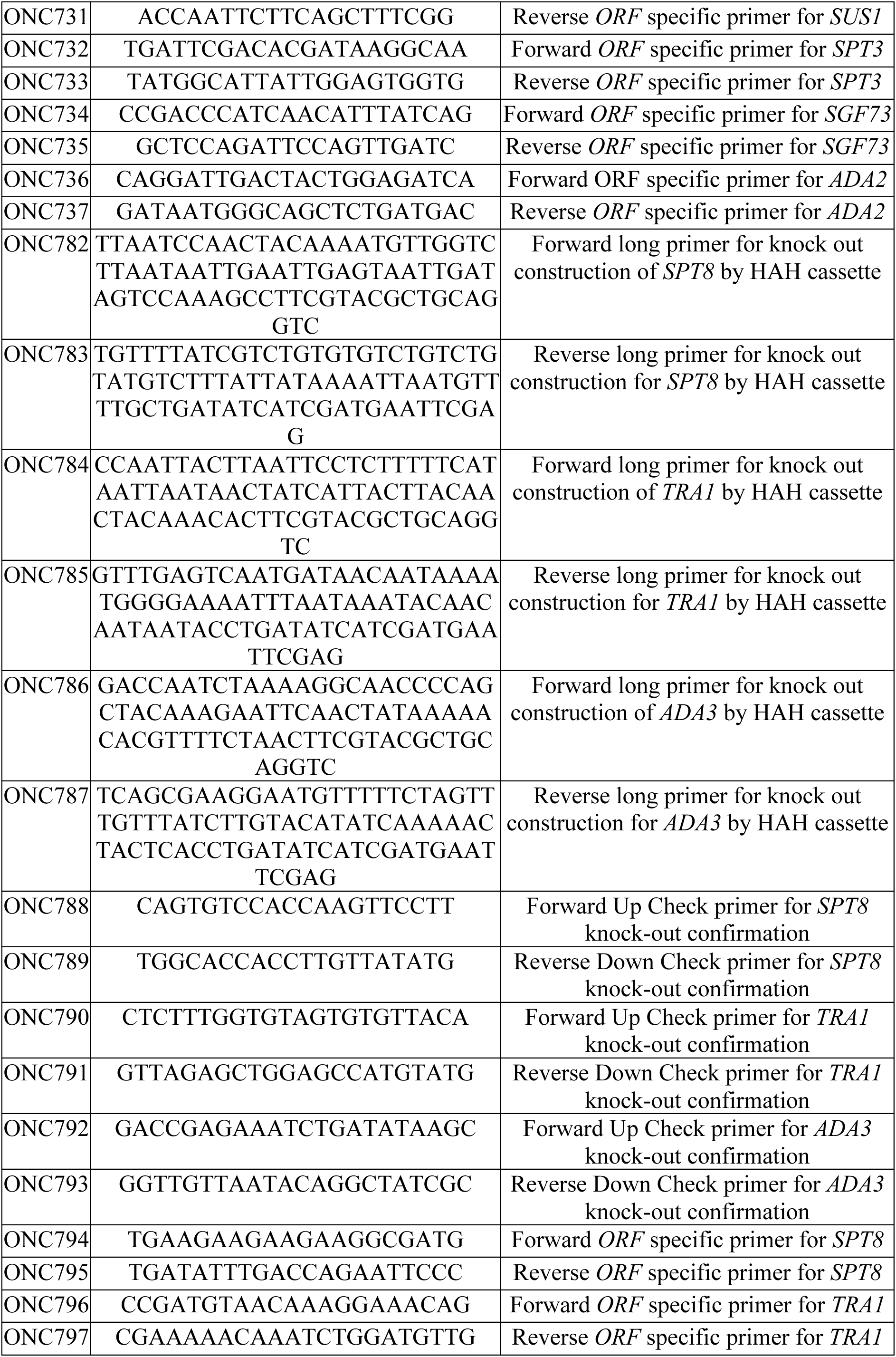
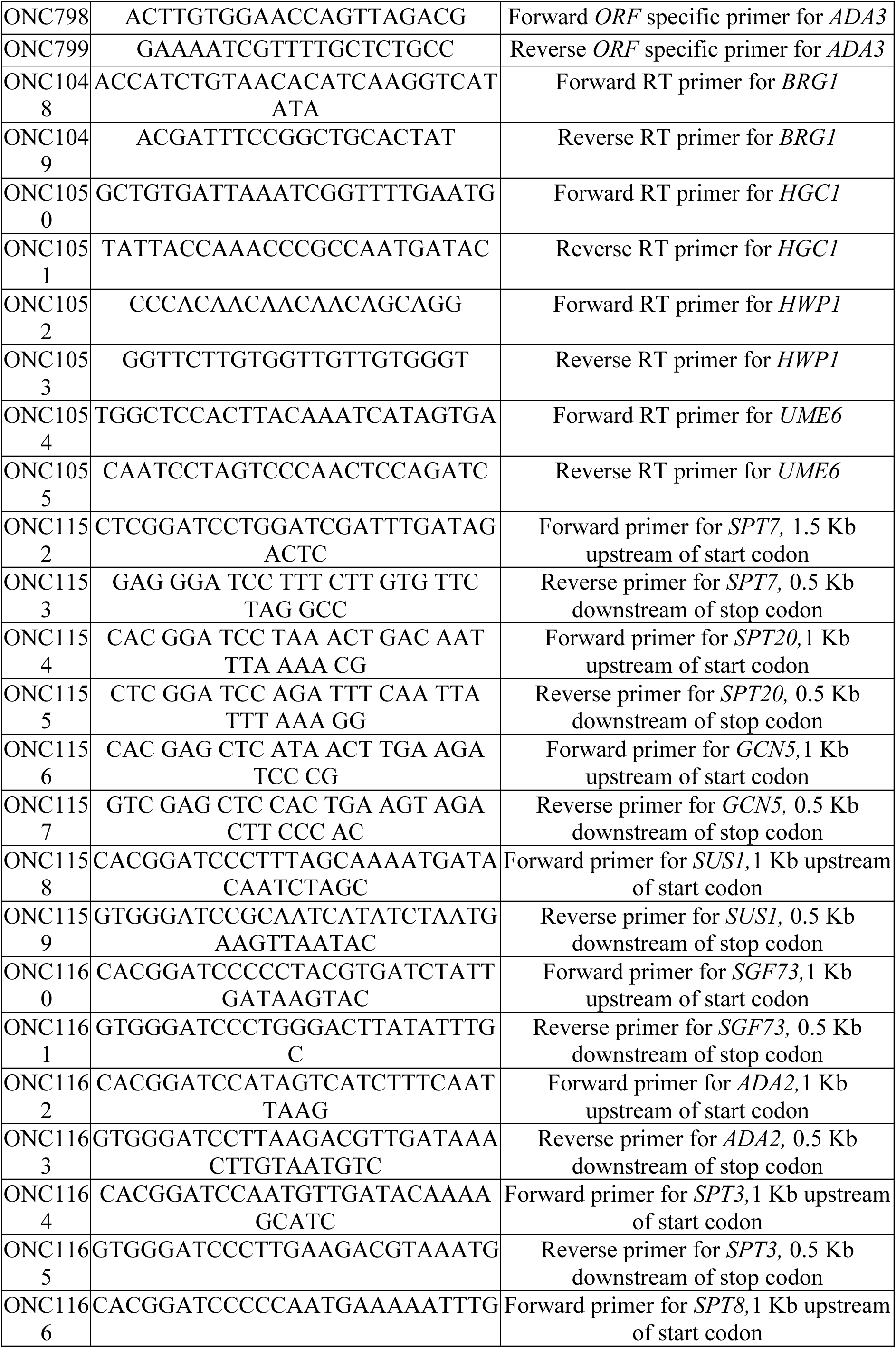
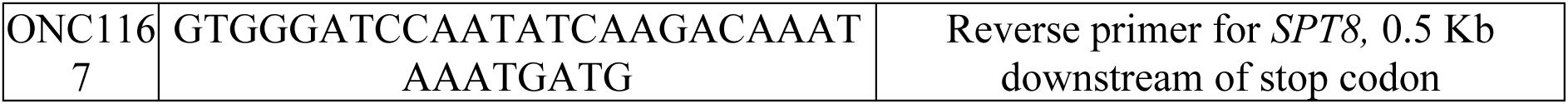
List of Oligonucleotides.

## Notes

### Competing Interest Statement

The authors have declared no competing interest.

