## Supplemental Figures for "Modular Architecture of the SAGA Complex Governs Stress Adaptation, Morphogenesis, and Histone Acetylation in *Candida albicans*"

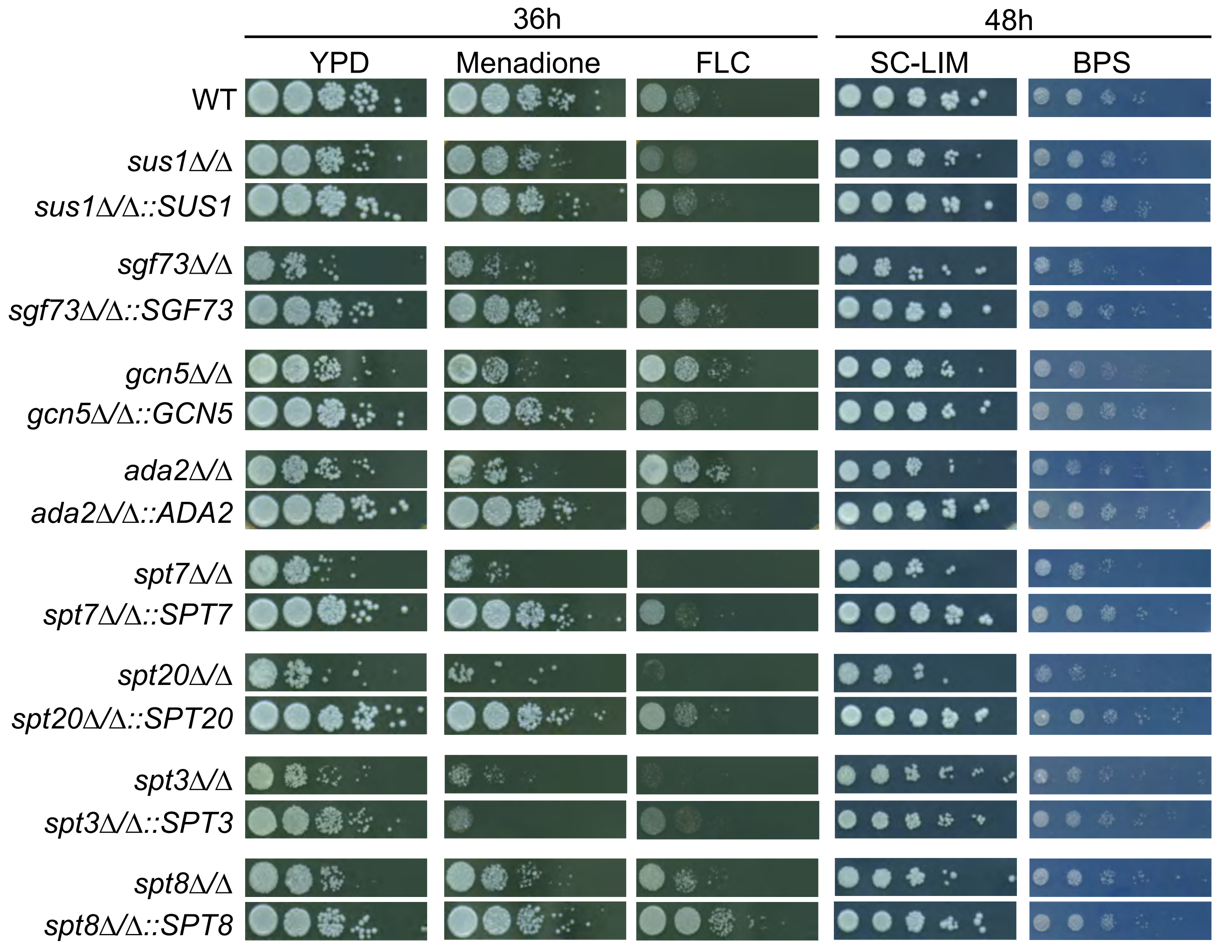

36h
48h
YPD
Menadione
FLC
SC-LIM
BPS
WT
sus1Δ/Δ
sus1Δ/Δ::SUS1
sgf73Δ/Δ
sgf73Δ/Δ::SGF73
gcn5Δ/Δ
gcn5Δ/Δ::GCN5
ada2Δ/Δ
ada2Δ/Δ::ADA2
spt7Δ/Δ
spt7Δ/Δ::SPT7
spt20Δ/Δ
spt20Δ/Δ::SPT20
spt3Δ/Δ
spt3Δ/Δ::SPT3
spt8Δ/Δ
spt8Δ/Δ::SPT8


**Figure S1**

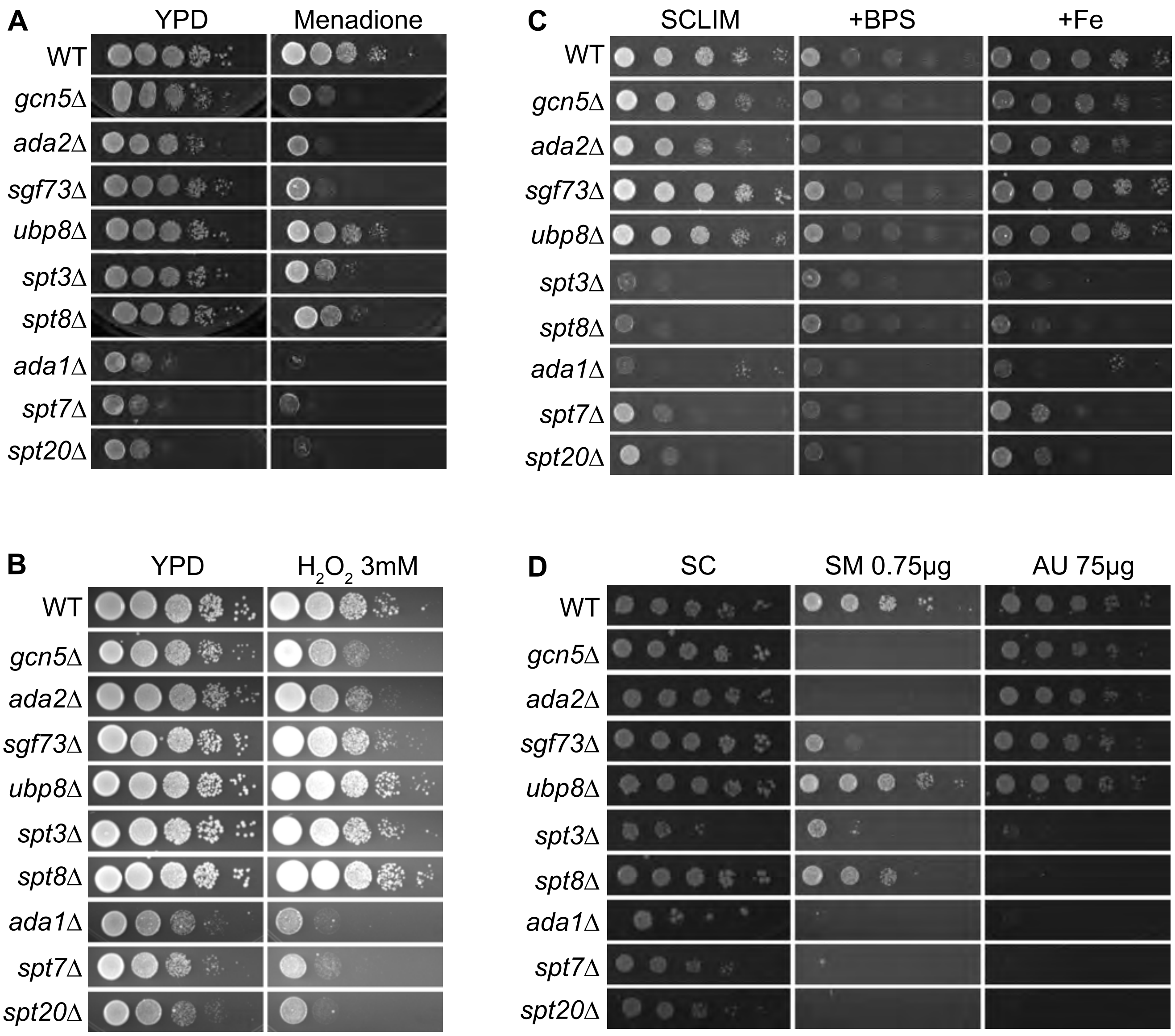


**E**

| Mutants/ Conditions | Ratio of Mutant/WT | | | | | |
|---|---|---|---|---|---|---|
| | Control | Menadione | $H_2O_2$ | BPS | SM | AU |
| *ubp8Δ* | 1.00 | 1.00 | 1.00 | 1.00 | 1.00 | 1.00 |
| *sgf73Δ* | 1.00 | 0.50 | 1.00 | 1.00 | 0.50 | 1.00 |
| *gcn5Δ* | 0.90 | 0.50 | 0.75 | 0.67 | 0.03 | 0.80 |
| *ada2Δ* | 0.90 | 0.50 | 0.75 | 0.67 | 0.03 | 0.80 |
| *spt7Δ* | 0.60 | 0.13 | 0.25 | 0.44 | 0.03 | 0.02 |
| *spt20Δ* | 0.60 | 0.13 | 0.25 | 0.44 | 0.03 | 0.02 |
| *spt3Δ* | 1.00 | 0.50 | 1.00 | 0.67 | 0.25 | 0.02 |
| *spt8Δ* | 1.00 | 0.50 | 1.00 | 0.67 | 0.75 | 0.02 |
| *ada1Δ* | 0.60 | 0.13 | 0.25 | 0.44 | 0.03 | 0.02 |

**Figure S2**

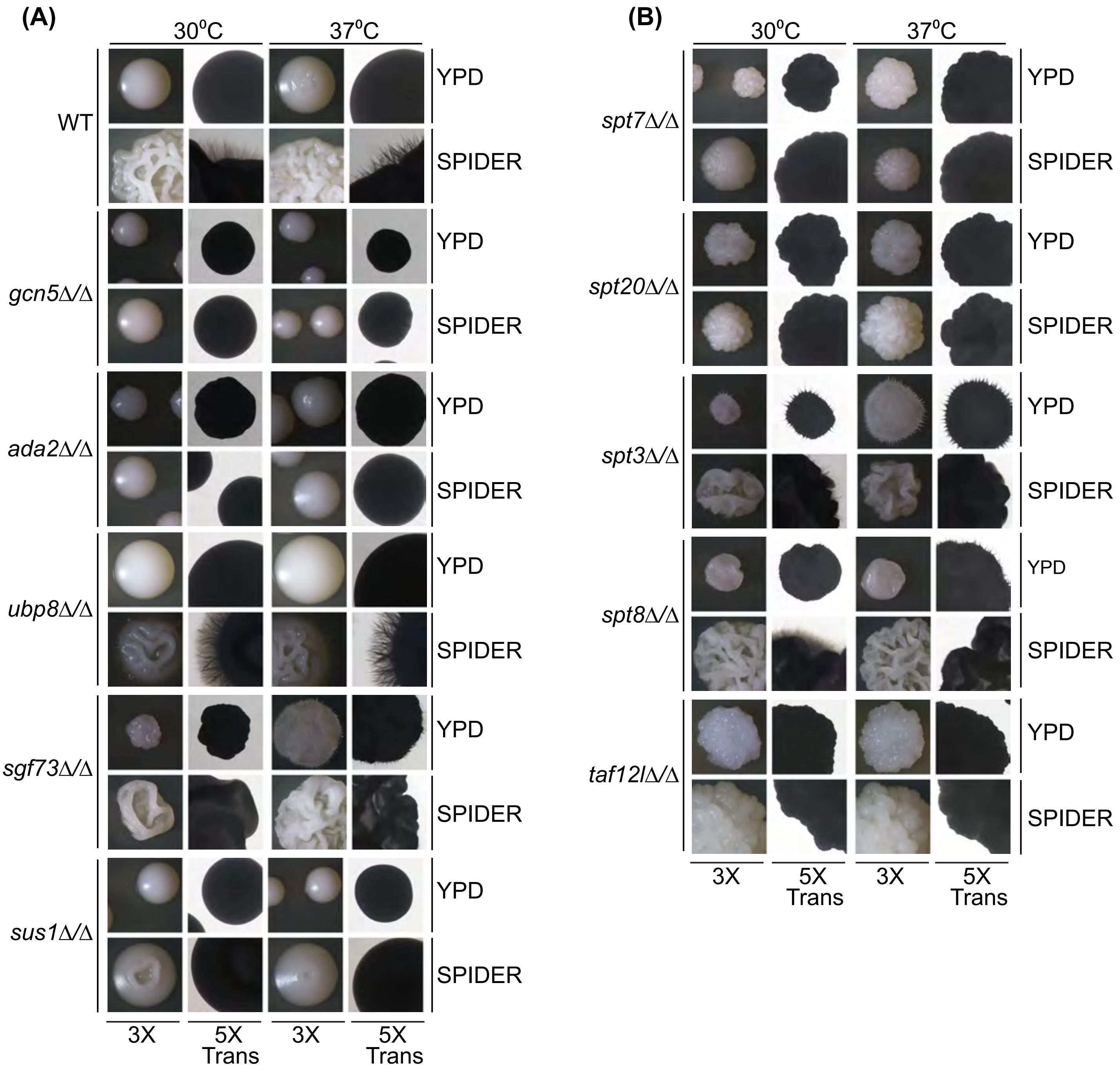
(A)
30°C
37°C
WT
gcn5Δ/Δ
ada2Δ/Δ
ubp8Δ/Δ
sgf73Δ/Δ
sus1Δ/Δ
YPD
SPIDER
3X
5X Trans
(B)
spt7Δ/Δ
spt20Δ/Δ
spt3Δ/Δ
spt8Δ/Δ
taf12lΔ/Δ

**Figure S3**

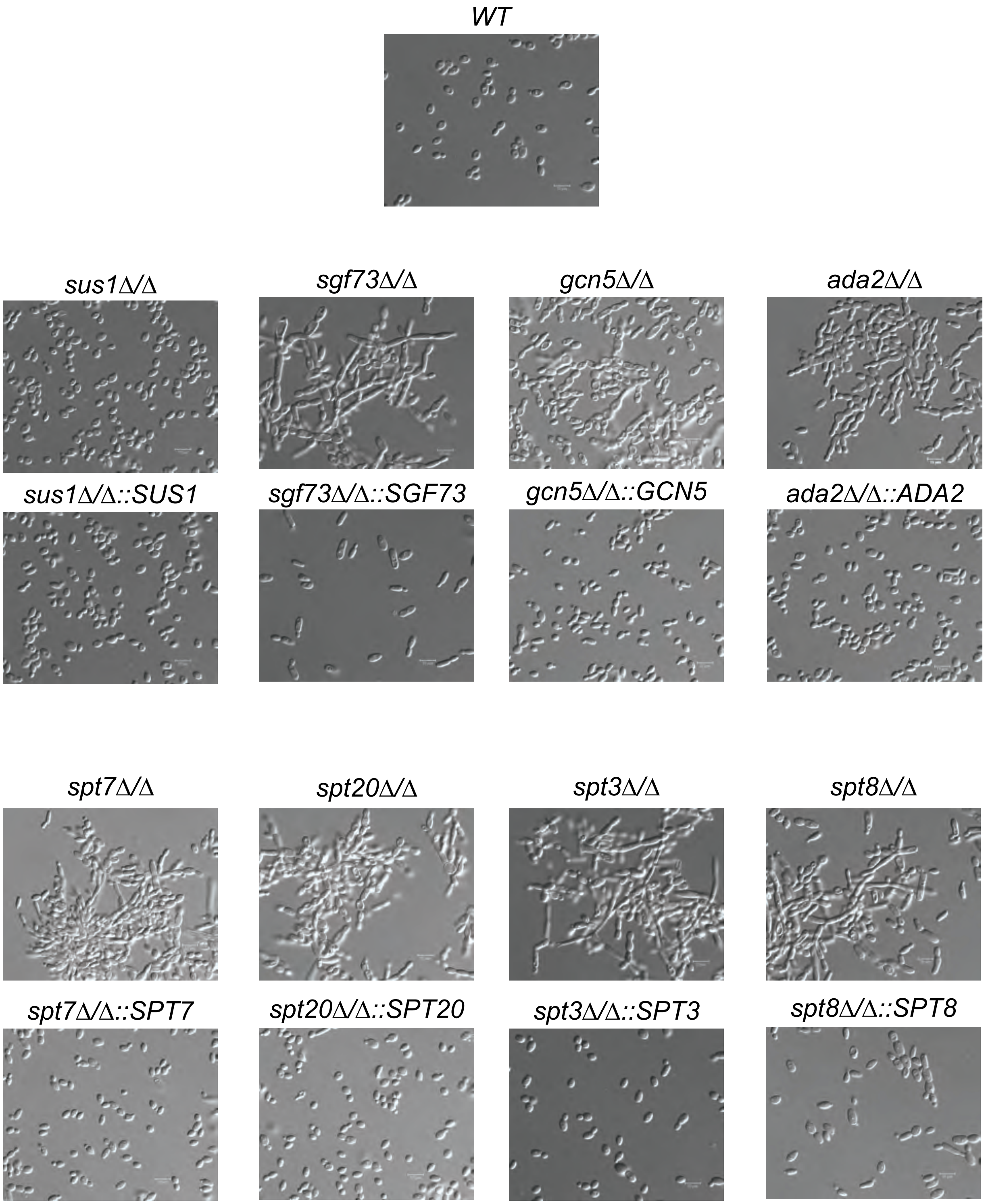

*WT*
*sus1Δ/Δ*
*sgf73Δ/Δ*
*gcn5Δ/Δ*
*ada2Δ/Δ*
*sus1Δ/Δ::SUS1*
*sgf73Δ/Δ::SGF73*
*gcn5Δ/Δ::GCN5*
*ada2Δ/Δ::ADA2*
*spt7Δ/Δ*
*spt20Δ/Δ*
*spt3Δ/Δ*
*spt8Δ/Δ*
*spt7Δ/Δ::SPT7*
*spt20Δ/Δ::SPT20*
*spt3Δ/Δ::SPT3*
*spt8Δ/Δ::SPT8*


**Figure S4**
